# A conserved activity for cohesin in bridging DNA molecules

**DOI:** 10.1101/757286

**Authors:** Pilar Gutierrez-Escribano, Matthew D. Newton, Aida Llauró, Jonas Huber, Loredana Tanasie, Joseph Davy, Isabel Aly, Ricardo Aramayo, Alex Montoya, Holger Kramer, Johannes Stigler, David S. Rueda, Luis Aragon

## Abstract

Essential processes such as accurate chromosome segregation, regulation of gene expression and DNA repair rely on protein-mediated DNA tethering. Sister chromatid cohesion requires the SMC complex cohesin to act as a protein linker that holds replicated chromatids together (*1, 2*). The molecular mechanism by which cohesins hold sister chromatids has remained controversial. Here, we used a single molecule approach to visualise the activity of cohesin complexes as they hold DNA molecules. We describe a DNA bridging activity that requires ATP and is conserved from yeast to human cohesin. We show that cohesin can form two distinct classes of bridges at physiological conditions, a “permanent bridge” able to resists high force (over 80pN) and a “reversible bridge” that breaks at lower forces (5-40pN). Both classes of bridges require Scc2/Scc4 in addition to ATP. We demonstrate that bridge formation requires physical proximity of the DNA segments to be tethered and show that “permanent” cohesin bridges can move between two DNA molecules but cannot be removed from DNA when they occur in *cis*. This suggests that separate physical compartments in cohesin molecules are involved in the bridge. Finally, we show that cohesin tetramers, unlike condensin, cannot compact linear DNA molecules against low force, demonstrating that the core activity of cohesin tetramers is bridging DNA rather than compacting it. Our findings carry important implications for the understanding of the basic mechanisms behind cohesin-dependent establishment of sister chromatid cohesion and chromosome architecture.

## Introduction

The establishment of sister chromatid cohesion is essential for accurate chromosome segregation during the mitotic cell cycle. Cohesin is an ATPase complex of the SMC (<u>s</u>tructural <u>m</u>aintenance of <u>c</u>hromosomes) family originally identified for its role in tethering sister chromatids from S phase until anaphase (*1, 2*). In addition to its function in sister chromatid cohesion, cohesin modulates the organisation of interphase nuclei and mitotic chromosomes (*1, 3, 4*). Studies in vertebrates have shown that cohesin complexes maintain contacts between different loci in *cis* and this way contribute to the folding of individual chromatids into distinct loops that provide an integral level of genome architecture (*1, 3, 4*). The current model for how SMC complexes, including cohesin, might form DNA loops involves the capture and bending of DNA segments followed by progressive enlargement of these to form loops (*5, 6*); this activity has been termed “loop extrusion”. Evidence for this model has been obtained from *in vitro* analysis of purified yeast condensin (*7*). Cohesin’s most prominent function is the tethering of sister chromatids, which is expected to involve an ability to bridge two DNA molecules in *trans*. It is currently not clear whether cohesin has a loop extrusion activity and if so how this might be linked to tethering of replicated chromatids. Mechanistically, we only have a vague idea of how cohesins might generate intermolecular tethers while mediating sister chromatid cohesion. Two main models have been proposed to explain cohesin function in sister chromatid cohesion: the “ring” or “embrace” model (*8, 9*), in which a single cohesin ring entraps both sister DNA molecules (*8*); and the “handcuff model”, whereby sister chromatid cohesion is mediated by the entrapment of sister DNAs in different cohesin complexes which interact with each other (*1, 10, 11*). The capture of the pair of dsDNA molecules during the establishment of sister chromatid cohesion by a single cohesin molecule in the “embrace model” has been proposed to occur by either, passage of the replisomes through the ring lumen of a DNA-bound cohesin or, (ii) when a DNA-bound cohesin captures a single-stranded DNA (ssDNA) at the fork which is then converted to dsDNA by DNA synthesis (*12*). Although cohesin complexes have been purified from fission yeast (*13*), frogs (*14*) and human cells (*15*), single molecule analyses of DNA bridging activities have not been reported. Purified cohesin complexes have been shown to exhibit DNA binding activity in a salt resistant manner (*16*) and to rapidly diffuse on DNA (*13-15*), however difussion was shown to be independent of ATP (*13-15*), suggesting that it is not at the core of its ATP-dependent activity.

Single molecule studies of purified yeast condensin have shown that this SMC complex compacts DNA molecules on magnetic tweezers (*17*), translocates directedly along linear DNA molecules in an ATP-dependent manner (*18*) and forms DNA loop-like structures on surfaced-tethered, flow-stretched DNA (*7*). Furthermore, while purified condensin exhibits robust ATPase activity in the presence of DNA (*17*), purified yeast cohesin is a poor ATPase (*19, 20*). Recent work has shown that the Scc2-Scc4 loader complex greatly stimulates cohesin’s ATPase activity (*19, 20*). Based on these findings we sought to investigate activities of purified budding yeast cohesin in the presence of the Scc2-Scc4 loader complex using two complementary single molecule approaches: DNA curtains and optical tweezers.

## Results

We purified budding yeast cohesin tetramers, containing Smc1, Smc3, Mcd1/Scc1, and Scc3, from exponentially growing yeast cultures (Supplementary Fig. 1A). Cohesin subunits were overexpressed in high-copy plasmids using galactose (*GAL*) inducible promoters. Purified material was obtained via affinity chromatography, using a triple StrepII tag fused to the Smc1 subunit, followed by passage through a HiTrap Heparin HP column (Supplementary Fig. 1A and Supplementary Table 1). Analysis of purified complexes by negative stain electron microscopy confirmed the presence of rod-shaped cohesin holocomplexes, the majority in a folded conformation (*21*) (Supplementary Fig. 1B). The Scc2-Scc4 complex was also purified from budding yeast (Supplementary Fig. 2A), using a similar strategy and exhibited DNA binding activity as expected (Supplementary Fig. 2B) (*19, 20*). Purified cohesin also bound plasmid DNA in a salt resistant manner consistent with the topological binding mode proposed for this complex (Supplementary Fig. 3A-B) (*16, 20*). Finally, we confirmed that our purified Scc2-Scc4 complex was able to stimulate cohesin ATPase activity (Supplementary Fig. 4) (*19, 20*).

Next, we sought to test whether budding yeast cohesin showed the behaviour described for cohesin purified from other organisms on DNA curtains (*13-15*). λ-DNA molecules (48.5 kb) were anchored to a lipid bilayer in a flow-cell surface and aligned into double-tethered DNA curtains using nano-fabricated barriers (*13*) (Fig. 1A). Quantum dots (Qdots) conjugated to antibodies against the hemagglutinin tag (HA3) fused to the C-terminal region of the Mcd1 kleisin subunit were used to visualise the complexes on DNA (Fig. 1B). On flowing the labelled cohesin complex over the DNA curtains binding was observed at low ionic strength (Supplementary Fig. 5A). The chamber was flushed with a high ionic strength buffer to remove non-topologically bound complexes (Supplementary Fig. 5A). While a large fraction of cohesin complexes dissociated, bound complexes diffused along the DNA (Supplementary Fig. 5B). The binding preference of cohesin to more A/T rich regions reported earlier (*13*) was also observed (Supplementary Fig. 5C-E). The diffusion coefficients correlated with the ionic strength of the buffer (Supplementary Fig. 5F). The survival probabilities of cohesin were not affected by the addition of ATP, or the ATP analogues ADP and ATP*γ*S (Fig. 1C). We found that the presence of Scc2-Scc4 enhanced the ability of cohesin to stay bound on the DNA (Fig. 1D), however the presence of nucleotides did not alter cohesin stability (Fig. 1D). Therefore, these results are consistent with the activities observed for cohesin from other organisms (*13, 15*) and show that budding yeast cohesin can diffuse on DNA curtains in an ATP independent manner.

**Figure 1.**
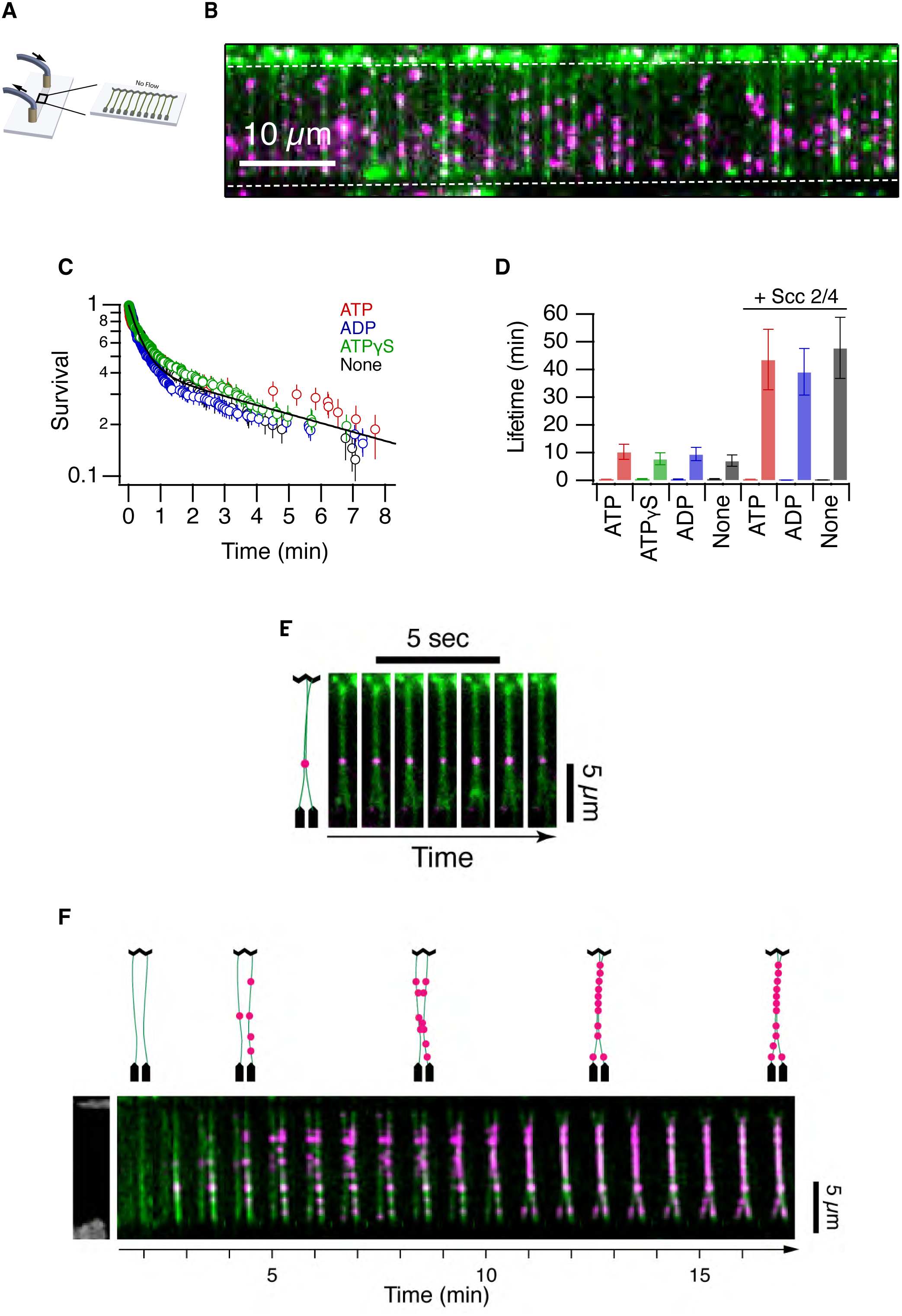
Analysis of yeast cohesin on DNA curtains. **A**. Schematic representation of double-tethered DNA curtains used in the study. **B**. Image of cohesin tagged with quantum dots (QD) (magenta) bound to λ-DNA stained with YOYO1 (green). Scale bar 10 μm. **C**. Survival probability plots of cohesin in the presence of ATP, ADP, ATP*γ*S or no nucleotide. **D**. Lifetimes of cohesin (fast phase and slow phase) in the presence/absence of Scc2-Scc4 and different ATP analogues. Error bars are 68% confidence intervals from bootstrapping. **E.** Image of a pair of double-tethered DNA curtains bound by cohesin in the presence of ATP and Scc2-Scc4. DNA molecules are in green, and cohesin is in magenta. Scale bar 5μm. Diagrammatic representation is shown (left). **F.** Time lapse images of a pair of double-tethered DNA curtains bound by cohesins as they are tethered in the presence of ATP and Scc2-Scc4. DNA molecules are in green, and cohesin is in magenta. Scale bar 5μm. Diagrammatic representation is shown (top).

In our DNA curtain experiments we made an observation not reported in earlier studies (*13, 15*). Cohesin signals were often observed bound between what appeared to be two fused DNAs (Fig. 1E). DNA pairing events with cohesin labelling (Fig. 1F) formed under low salt conditions in the presence of ATP and Scc2-Scc4 complex (Fig. 1F), but persited when the chamber was flushed with a high ionic strength buffer, raising the possibility that topologically bound complexes mediated these events. To further explore this, we decided to use a dual-trap optical tweezer with confocal fluorescence microscopy capabilities. A similar approach has been previously used in the study of protein-DNA interactions (*22*). Briefly, we tether a *λ*-DNA molecule with biotinylated ends to two optically-trapped streptavidin-coated polystyrene beads, enabling us to accurately apply and measure forces on the captured DNA molecule. We performed our experiments in multi-channel laminar flow cells where we had the possibility to move the tethered DNA between different flow lanes containing distinct protein complexes and buffers. In addition, we were able to image the tethered DNAs using confocal fluorescence microscopy. Overall, the approach allows increased experimental control over DNA curtains. Proteins can be added, removed or incubated in different salt conditions sequentially and the physical effect of their activities can be measured accurately on a single DNA molecule.

To test for the formation of intramolecular cohesin bridges *in cis*, we adapted a previously published protocol that measures protein-mediated DNA bridging (*23, 24*) (Fig. 2A). First, we captured a single λ-DNA molecule and generated a force–extension (FE) curve in the absence of protein by extending the molecule slightly beyond its contour length (∼16 µm). We then moved the DNA to a channel containing 1 nM cohesin, 2.5nM Scc2-Scc4 complex and 1mM ATP in 50mN NaCl and incubated for 30s in a relaxed conformation (∼3 µm between beads). Following incubation, the relaxed DNA was then moved to a channel without protein but containing 1mM ATP in 125mM NaCl. Re-extending the DNA in the buffer channel yielded FE curves with sawtooth features at extensions shorter than the contour length (Fig. 2B; cohesin + Scc2/4). This is characteristic of intramolecular bridge-rupture events (*23, 24*) (Fig. 2A; right panel) and shows that cohesin can tether the DNA in *cis* forming a protein-mediated bridge between different segments of the molecule, thus creating an intramolecular loop. Importantly, when we repeated this protocol in the presence of 1nM cohesin and no Scc2-Scc4 (Fig. 2B; cohesin), or 2.5nM Scc2-Scc4 and no cohesin (Fig. 2B; Scc2/4), FE curves identical to those of the initial naked DNA were observed. This demonstrates that no protein-mediated bridges were formed (Fig. 2A; left panel). Similarly, incubating 1nM cohesin and 2.5nM Scc2-Scc4 complex in the absence of ATP, or with the ATP analogues, ADP or ATP*γ*S yielded FE curves identical to those of naked DNA (Supplementary Fig. 6A). To confirm the requirement of ATP, we repeated the protocol in the presence of 1nM cohesin ATPase mutant (K38I) (Supplementary Fig. 7), and 2.5nM Scc2-Scc4 (Fig. 2B; cohesinK38I + Scc2/4). FE curves identical to those of the naked DNA were observed (Fig. 2B; cohesinK38I + Scc2/4). Therefore, our results demonstrate that the DNA bridging activity requires ATP and depends on the Scc2-Scc4 loader complex.

**Figure 2.**
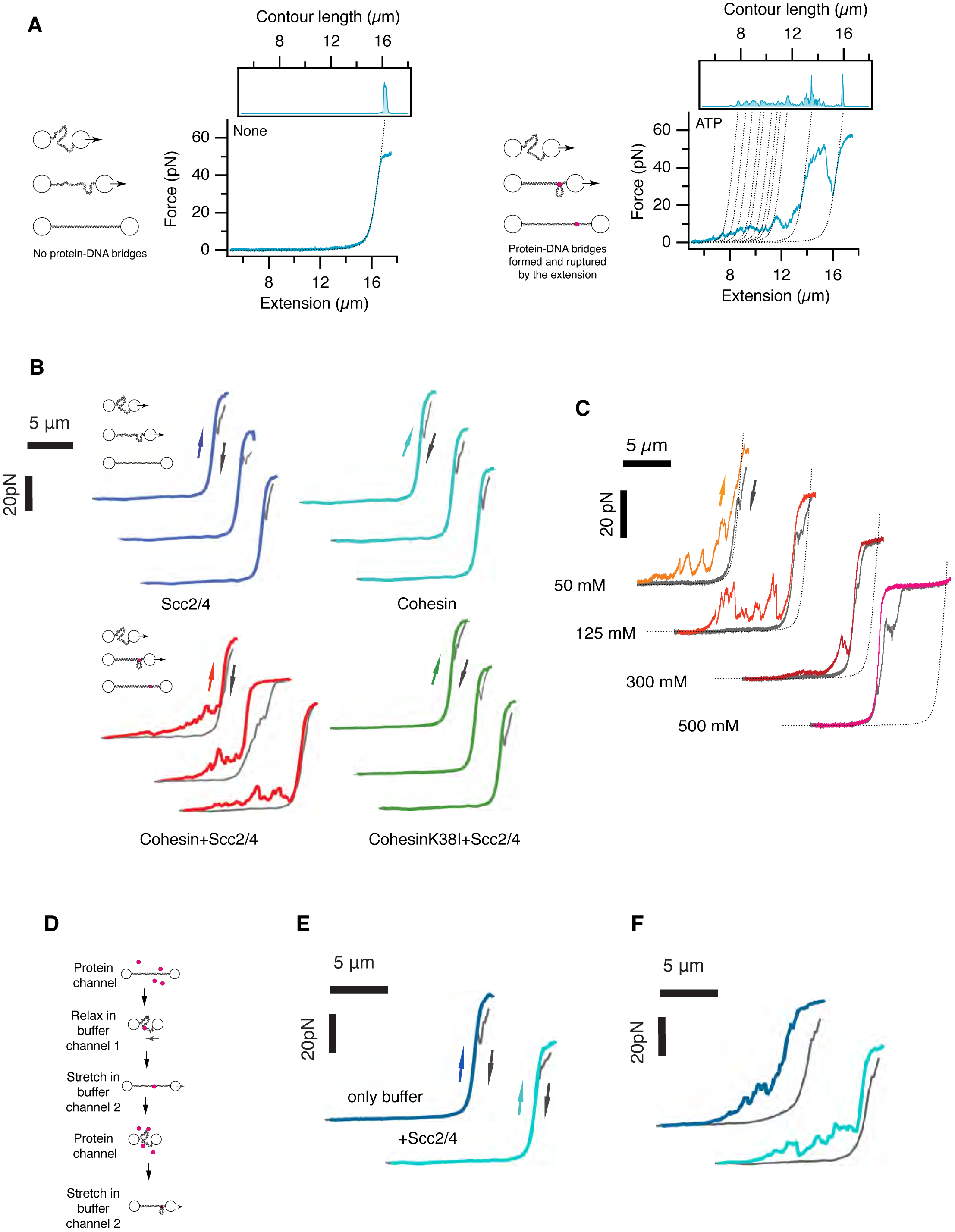
Cohesin bridges DNA in an ATP and Scc2-Scc4 dependent manner. **A**. Schematic representation of Force-Extension (FE) curves for λ-DNA exhibiting the presence (right diagramme and graph) and absence (left diagramme and graph) of protein-DNA bridges. Dotted line is fit to worm-like-chain (WLC) for naked DNA. **B**. FE curves for λ-DNAs pre-incubated with 1nM cohesin and 2.5nM complex and 1mM ATP (Cohesin+Scc2/4), 1nM cohesin and 1mM ATP (Cohesin), 2.5nM Scc2-Scc4 and 1mM ATP (Scc2/4) or 1nM cohesin ATPase mutant and 2.5nM complex and 1mM ATP (CohesinK38I + Scc2/4). Schematic diagram of the experimental design is shown on the left. After capturing a single DNA molecule between two optically trapped beads, DNA was incubated in the presence of protein (as indicated) in a relaxed conformation (3 μm bead distance) for 30s in 50mM NaCl and then moved to a buffer channel with 125mM NaCl for extension and measurements. Only incubation with 1nM cohesin and 2.5nM complex and 1mM ATP (Cohesin+Scc2/4) showed DNA bridge rupture events. **C.** FE curves in the presence of increasing ionic strength. High salt favours topologically constrained and permanent DNA bridges. **D** Schematic representation of the experimental design to test cohesin second DNA capture. After capture of λ-DNA between the two optically trapped beads, DNA is extended and incubated for 30 seconds in the protein channel. DNA is moved to a buffer channel, then relaxed (3μm bead distance) and incubated for 30 seconds before re-extension to test for DNA bridges (Graphs in ***E***). The extended DNA is then incubated in a relaxed position in the protein channel, then moved to buffer channel and extended to confirm that bridges can be formed when protein is loaded while DNA is relaxed (Graphs in ***F***). **E**. λ-DNA incubated with 1nM cohesin, 2.5nM Scc2-Scc4 complex and 1mM ATP in an extended conformation. Then moved to a buffer channel (125mM NaCl) in the presence of 1mM ATP (buffer only - dark blue) or 2.5nM Scc2-Scc4 complex and 1mM ATP (+Scc2/4 - light blue). DNAs were re-extended and the FE curves shown recorded. **F**. The λ-DNA molecules in (***E***) were further incubated in a relaxed position (3μm bead distance) in the presence of 1nM cohesin, 2.5nM Scc2-Scc4 complex and 1mM ATP DNAs. DNAs were moved to the buffer only channel (125mM NaCl containing 1mM ATP) and re-extended. FE curves show the presence of DNA bridge rupture events. Experiments involving FE curves were repeated a minimum of 5 times per condition.

Next, we tested the effect of ionic strength on cohesin bridging (Fig. 2C). Cohesin bridges were observed at all salt concentrations tested (Fig. 2C). The length of DNA extension released during the rupture of a DNA bridge can be directly related to the loop size encompassed by the bridge. We analyzed the sizes of the DNA bridges from the FE curves (Supplementary Fig. 6B) and found that the distribution of loop sizes is exponential with a characteristic size of ∼900bp, consistent with a model of random bridge formation (*5, 6*) (Supplementary Fig. 6B). Interestingly, most of the small sawtooth peaks observed at a low forces and extensions disappeared in high salt, while the overall contour length of the DNA remained reduced (Fig. 2C). We also recorded FE curves when we relaxed tethers (Fig. 2B-C; reverse arrows) after the extensions (Fig. 2B-C; forward arrows). These showed that compaction due to DNA bridges formed at low salt concentrations were lost after extension (Fig. 2C; reverse arrows; 50mM NaCl). However, relaxation of tethers with DNA bridges formed at high salt concentrations showed compaction events that had resisted after extension (Fig. 2C; reverse arrows; 300 and 500mM NaCl). These results demonstrate the existence of two distinct types of cohesin bridging events, (i) one predominantly occurring at low salt that is characterised by frequent interactions that are “reversible” and can be disrupted by moderate force (5-40pN) and, (ii) a second “permanent” bridge class that resists higher ionic strength conditions and full physical stretching of the DNA molecule. Both classes of DNA bridges were not observed when an ATPase mutant cohesin complex (SMC3-K38I) was used (Fig. 2B; cohesinK38I + Scc2/4) confirming that the ATPase activity of the complex is a requirement for both types of bridges. Importantly, in experiments done at physiological salt concentration (125mM NaCl) we detected both types of cohesin bridges (Fig. 2C; 125mM NaCl), raising the possibility that both bridge classes might occur *in vivo*. Next, we tested whether “permanent” bridges could resist repeated extensions. We performed two cycles of bead extension and relaxation and confirmed the persistence of the “permanent” cohesin bridge (Supplementary Fig. 8). We conclude that “permanent” cohesin bridges resist high stretching forces, and that the complexes mediating these tethers cannot be displaced from the DNA molecules. This explains the repeated detection of the same bridge characteristics on FE curves during the two cycles of bead extension and relaxation (Supplementary Fig. 8).

Recent studies using purified cohesin from *S. pombe* have shown that cohesin can capture a second DNA but only if single-stranded (*12*), suggesting that this event is likely to occur at replication forks (*12*). The second capture of the single stranded molecule was dependent on the presence of cohesin loader and ATP (*12*). Our results show that cohesin purified from *S. cerevisiae* is able to trap two dsDNA segments in the same λ-DNA molecule (Fig. 2B-C), however our tethering assay could not differentiate whether the two dsDNA segments are captured sequentially or in a single step, as we had incubated the DNA in a relaxed position (with the two DNA segments in proximity). To distinguish whether one or two events were involved in the formation of the cohesin tethers observed, we sought to test whether cohesin could capture a second DNA segment (within the same molecule) after initial loading. To this aim, we captured a single λ-DNA molecule and generated a force–extension (FE) curve. We maintained the DNA in an extended position (∼14 µm between beads) using a pulling force of 5pN (Fig. 2D) and loaded cohesin by moving the DNA to a channel containing 1nM cohesin, 2.5nM Scc2-Scc4 complex and 1mM ATP in 50mM NaCl. We incubated the DNA for 30s (Fig. 2D) before moving it to a different channel containing 1mM ATP in 50mM NaCl. We then relaxed the DNA conformation (∼3 µm between beads) to allow DNA segments to come into proximity (Fig. 2D) and incubated in the relaxed conformation for 30 additional seconds. The DNA was then moved to a different channel containing 1mM ATP in 125mM NaCl and extended to record an FE curve. The curve obtained after re-extension of the DNA was identical to the initial naked DNA profile (Fig. 2E; only buffer and Supplementary Fig. 9). We obtained a similar result when we included 2.5nM Scc2-Scc4 complex and 1mM ATP in the channel where we relaxed the DNA (Fig. 2E; +Scc2/4 and Supplementary Fig. 9). These results show that loaded cohesin is unable to capture a second DNA segment. To confirm that DNA bridges could be formed in the same DNA in one step, we relaxed the molecules used in the experiments and incubated them 30s in a channel containing 1nM cohesin, 2.5nM Scc2-Scc4 complex and 1mM ATP. When molecules were re-extended and the resulting FE curves recorded, they confirmed the formation of DNA bridges (Fig. 3F and Supplementary Fig. 9). In addition, we confirmed that cohesin complexes can bind to extended DNAs using a published DNA friction protocol (*25*) (Supplementary Fig. 10). Therefore, our results are consistent with previous reports (*12*) showing that cohesin bound to DNA cannot undergo a second capture event involving a dsDNA molecule. We conclude that cohesin establishes bridges between two dsDNAs in a single event which requires physical proximity of the DNA segments that are tethered. Importantly, previous studies could not evaluate the possibility that cohesin could capture two dsDNAs simultaneously (*12*).

**Figure 3.**
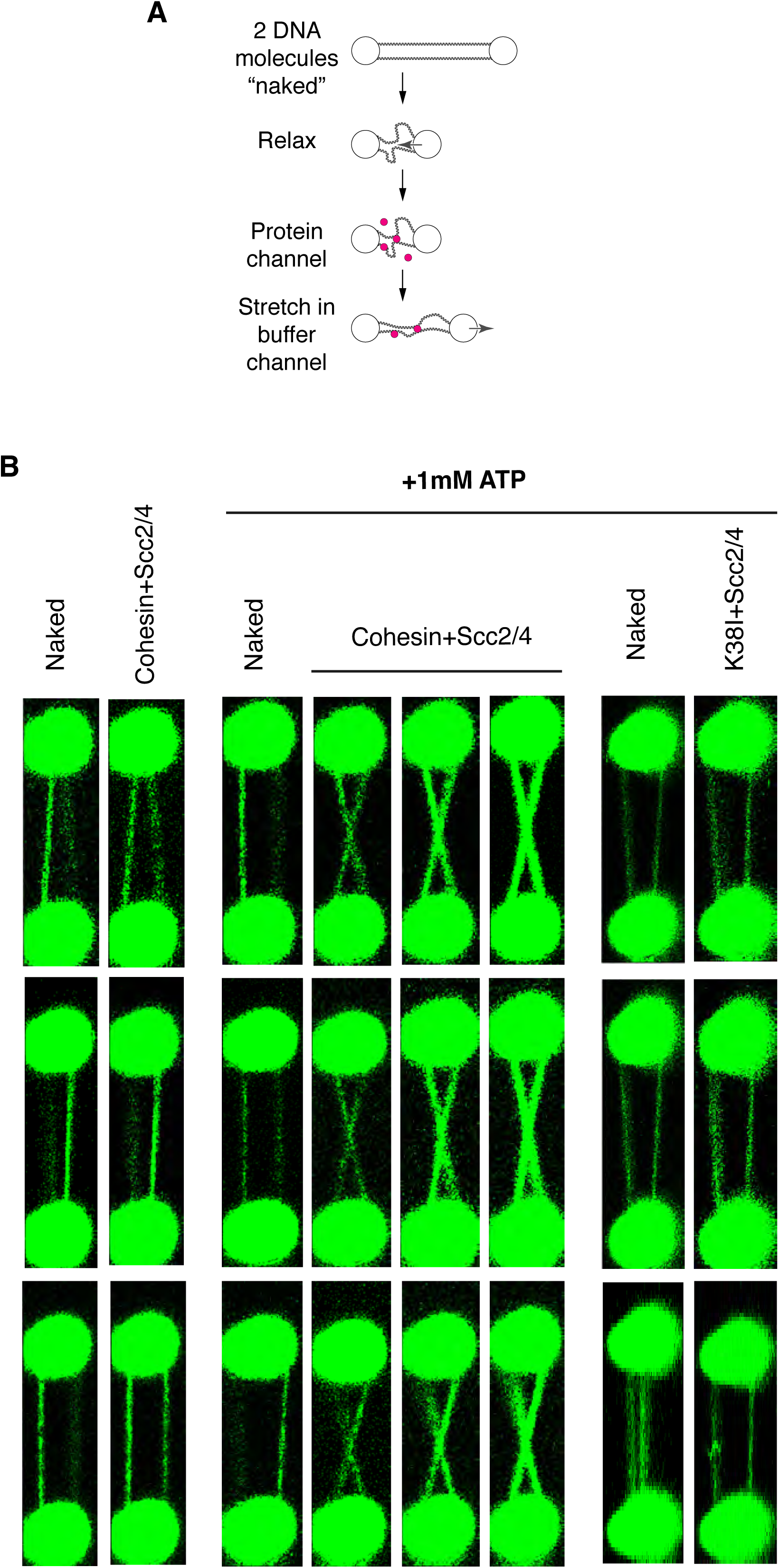
Cohesin bridges DNA molecules in *trans*. **A**. Schematic representation of the experimental design for the dual trap optical tweezer assay to generate permanent intermolecular cohesin bridges. Two λ-DNA molecules are tethered between the two beads and incubated in a relaxed position (3μm bead distance) in the presence/absence of protein in 50mM NaCl buffer. The relaxed molecules are then moved to a different channel containing 300mM NaCl and re-extended. Imaging is done before incubations and after re-extension in the 300mM NaCl buffer using 50nM of SYTOX Orange to visualise DNA. **B.** Two λ-DNA molecules were tethered and treated as described in (***A***) and incubated with either (i) 1nM cohesin, 2.5nM Scc2-Scc4 and no ATP (Cohesin+Scc2/4 - left), (ii) 1nM cohesin, 2.5nM Scc2-Scc4 and 1mM ATP (Cohesin+Scc2/4 - middle), or (iii) 1nM cohesin ATPase mutant K38I, 2.5nM Scc2-Scc4 and 1mM ATP (K38I+Scc2/4 - right). Imaging was performed before incubation and after DNA re-extension in 300mM NaCl buffer, to minimise DNA entanglement. Images from three independent experiments for each category are shown. Bridging experiments were repeated a minimum of 6 times.

Next, we investigated whether cohesin could form intermolecular bridges. We developed an intermolecular bridging assay, where two dsDNA molecules are tethered in parallel between the pair of beads, and tested the ability of cohesin to form bridges between these two molecules (Fig. 3A). After confirming the presence of two DNA molecules tethered in parallel between the beads using Sytox Orange (Fig. 3B; Naked) the DNA was incubated in a relaxed state to bring the DNAs into proximity (∼3 µm bead distance) in the presence of 1nM cohesin, 2.5 nM Scc2-Scc4 and 1mM ATP in 50mM NaCl for 30s. Then, the DNAs were moved to a buffer-only channel (300 mM NaCl and 1mM ATP). Strikingly, clear bridging was observed between the two molecules on re-extension (Fig. 3B; Cohesin + Scc2/4 +1mM ATP). DNA bridges did not form in the absence of ATP (Fig. 3B; Cohesin + Scc2/4) or when we used cohesin ATPase mutant complex (Fig. 3B; K38I+Scc2/4 +1mM ATP), confirming that cohesin’s ATPse activity is required. Bridge formation in this assay was very efficient; out of 10 molecules tested, 8 showed intermolecular bridges and 2 showed intramolecular bridging on the two individual DNAs. Intermolecular bridges always appeared to be near the midpoint of the DNA (Fig. 3B; Cohesin + Scc2/4). Potential reasons to explain this include the fact that the central region of λ-DNA molecules is rich in A/T content where cohesin might bind preferentially. Alternatively, cohesin might be able to slide on the DNA while maintaining tethers and therefore move to the centre regions as the molecules are extended. To further characterise this, we used a quadruple-trap optical tweezer setup which allows the independent manipulation of the two DNA molecules (*25*).

We first captured two single λ-DNA molecules using a pair of traps for each (DNA1 between traps 1-2 and DNA2 between traps 3-4) in a parallel conformation (Supplementary Fig. 11). Both DNA molecules were stretched close to their contour lengths (∼16 µm). We then manipulated DNA2 using beads 3 and 4 and moved it upwards (in the *z*-direction) before rotating it 90 degrees and moving it into a crossed conformation directly above DNA1 (Supplementary Fig. 11). We then lowered DNA2 to its original z-position and relaxed it to ensure physical contact between the two DNA molecules at the junction point (Supplementary Fig. 11). We then moved the crossed DNAs into a different channel containing 1nM cohesin, 2.5 nM Scc2-Scc4 and 1mM ATP in 50 mM NaCl and incubated it for 60s before returning the DNAs to a channel containing 1mM ATP in 300 mM NaCl. We reversed the manipulation of DNA2, first moving bead 3 upwards and over DNA1 before manipulating beads 3 and 4 so that DNA2 was rotated −90 degrees and lowered back to the original position where DNA1 and DNA2 were parallel to each other. That configuration was then moved to a channel containing Sytox Orange to visualise the DNA molecules. We observed that DNA1 and DNA2 were bridged (Supplementary Fig. 11), as expected from our analysis of parallel DNA bridging in the dual trap optical tweezer setup (Fig. 3B; Cohesin + Scc2/4 +1mM ATP). We then tested whether moving DNA2 using simultaneous movement of beads 3 and 4 in the x axis would cause the sliding of the bridge along DNA1 (Fig. 4A). Indeed, we observed that the bridge could be moved showing that cohesin can slide on DNAs while tethering two DNA molecules in *trans* (Fig. 4A). Importantly, when we applied force to disrupt the bridge (moving bead 3 down in the y axis; i.e. away from beads 1 and 2) (Fig. 4B) we could not break apart the cohesin tether. At high forces, the interaction between the ends of the DNAs and the beads snapped (Fig. 4B). Amazingly, cohesin bridges resisted this and half of DNA2 could be observed hanging from the bridge (Fig. 4B). We conclude that “permanent” intermolecular cohesin bridges can slide on DNA and resist high force.

**Figure 4.**
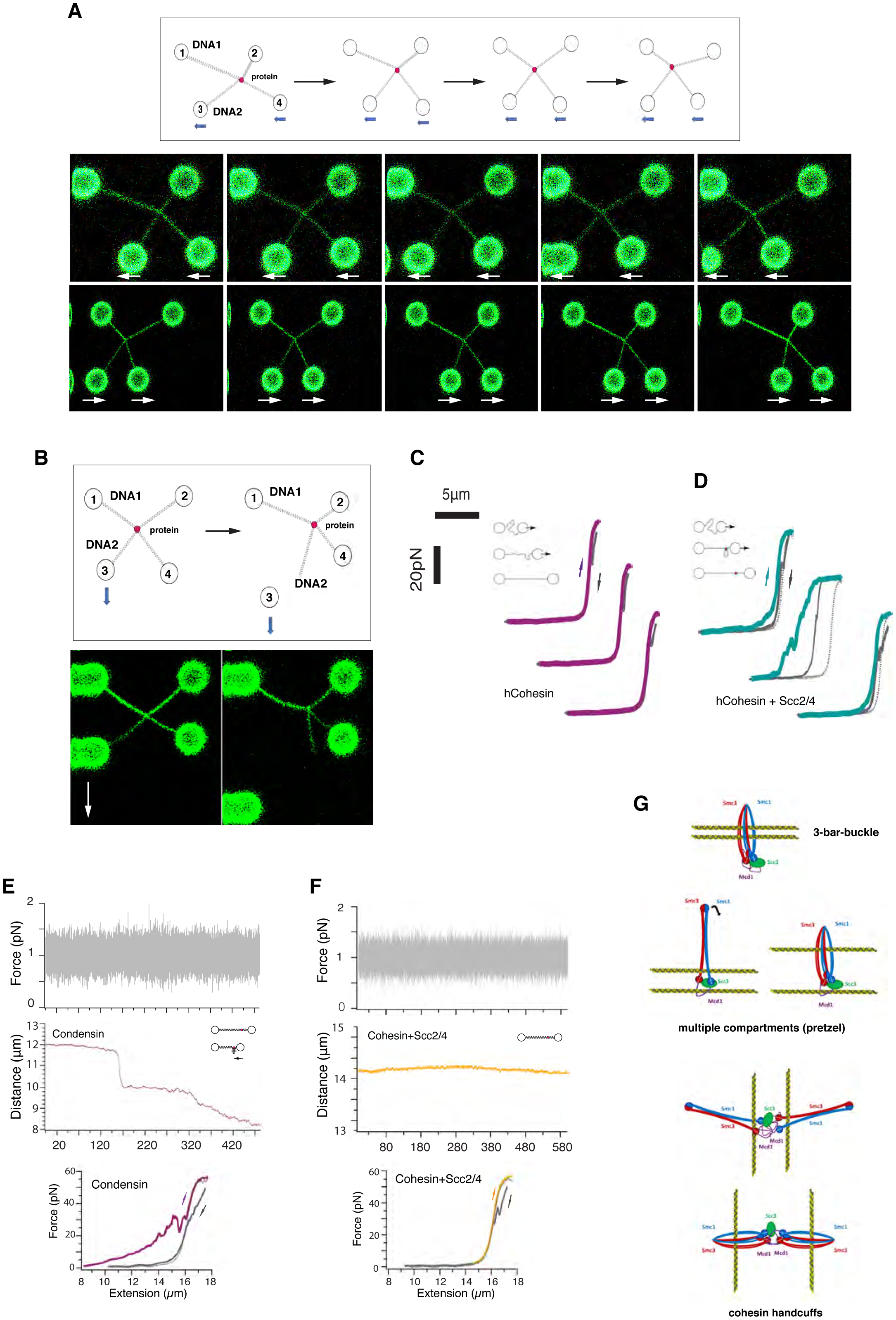
Intermolecular cohesin bridges slide on DNA. **A.** Schematic representation of the experimental design to test for sliding of permanent cohesin bridges (top diagramme). Following the formation of an intermolecular cohesin bridge (see Supplementary Fig. 11 for details in bridge formation protocol). Beads 3 and 4 are moved together in the x axis to slide the bridge along DNA1. Images showing two representative sliding experiments are shown. Experiments were performed in a buffer containing 300mM NaCl and 50nM of SYTOX Orange. The experiment was performed three times and sliding was observed in all cases. **B.** Schematic representation of the experimental design to disrupt intermolecular cohesin bridges. Following the formation of an intermolecular cohesin bridge, beads 3 is moved down in the y axis until one of the DNA ends losses contact with the bead. Imaging was performed before and after the pull in a buffer containing 300mM NaCl and 50nM of SYTOX Orange. Representative experiment is shown. The experiment was repeated 3 times. **C.** Force-Extension (FE) curve for λ-DNA pre-incubated with 1nM human Cohesin and 1mM ATP in 125mM NaCl buffer (hCohesin). Dotted line is fit to worm-like-chain (WLC) model. After capturing a single DNA molecule between two optically trapped beads, DNA was incubated in the presence of protein in 50mM NaCl buffer in a relaxed conformation (3 μm bead distance) for 30s and then moved to the 125mM NaCl buffer channel for extension and measurements. No evidence of DNA bridges is observed in this condition. Experiments involving FE curves were repeated 3 times. **D.** Force-Extension (FE) curve for λ-DNA pre-incubated with 1nM human Cohesin, 2.5nM yeast Scc2-Scc4 and 1mM ATP in 125mM NaCl buffer (hCohesin + Scc2/4). Experimental procedure as in (***C***). FE curves exhibit multiple rupture events indicating the presence of “reversible” and “permanent” DNA bridges. Experiments involving FE curves were repeated 3 times. **E.** DNA compaction trace for λ-DNA molecule extended using a force of 1pN (top). The DNA was tethered between two beads. One bead was clamped (fixed) while a 5pN force was applied to the second bead to maintain the molecule extended. The DNA was then incubated in the presence of 1nM condensin (1mM ATP in 50mM NaCl) (-condensin - magenta traces). Extended DNAs were then moved to a different channel containing 1mM ATP in 50mM NaCl and the extension force was reduced to 1pN. The distance between the beads was recorded over time. The FE curve for the λ-DNA full extension after incubation is shown (bottom). Additional examples can be found in Supplementary Fig. 15. Experiments were repeated 3 independent times. For graphical representation, force data were downsampled to 100Hz. **F.** DNA compaction trace for λ-DNA molecule extended using a force of 1pN (top) in the presence of 1nM Cohesin and 2.5nM Scc2-Scc4 complex (1mM ATP in 50mM NaCl) (right-yellow trace). Experimental procedure was as in (***E***). The distance between the beads was recorded over time. The FE curve for the λ-DNA full extension after incubation is shown (bottom). Additional examples can be found in Supplementary Fig. 15. Experiments were repeated 3 independent times. For graphical representation, force data were downsampled to 100Hz. **G.** Tentative models for cohesin bridging activity. Permanent DNA bridges slide when force is applied (***A***), however, when they occur in *cis*, cohesin complexes cannot slide off the DNA molecules (Fig. 2C and Supplementary Fig. 8). This demonstrates that the two DNA molecules (or DNA segments in the same molecule) tethered are located in different physical compartments within the protein. Either two compartments within one cohesin tetramer; the 3-bar-buckle and multiple compartment (pretzel) models, or different compartments of two cohesin complexes; the handcuff model.

Previous studies using purified cohesin did not report DNA bridging activities (*13-15*), however the studies did not employ budding yeast cohesin. We therefore decided to test whether the bridging activity observed is specific for *S. cerevisiae* cohesin tetramers or it has been conserved in cohesin from other organisms. To this aim, we purified the human cohesin (hCohesin) tetramer complex, containing hSmc1, hSmc3, hRad21 and Stag1 as described previously (*26*) (Supplementary Fig. 12). We then tested whether hCohesin could bridge DNA intramolecularly. We captured a single λ-DNA molecule and generated a FE curve in the absence of protein to confirm the presence of naked DNA. We then moved the DNA to a channel containing 1nM hcCohesin and 1mM ATP in 50mN NaCl and incubated it for 30s in a relaxed conformation (∼3 µm between beads). We then moved the relaxed DNA to a channel without protein in the presence of 1mM ATP in 125mM NaCl. Re-extending the DNA resulted in FE curves with a naked DNA profile (Fig. 4C; hCohesin), demonstrating that hCohesin cannot promote DNA bridges. Although we could not obtain hScc2-Scc4, we decided to test whether budding yeast loader complex Scc2-Scc4 (scScc2-Scc4) had any effect on hCohesin activity. To this aim we repeated the intramolecular DNA bridging assays with hCohesin and included Scc2-Scc4 loader complex in the incubations. Relaxed DNA was incubated in the presence of 1nM hCohesin tetramer, 2.5nM scScc2-Scc4 complex and 1mM ATP in 50mM NaCl. The relaxed DNA was then moved to a channel with 1mM ATP in 125mM NaCl. Re-extension yielded the sawtooth features characteristic of intramolecular bridge-rupture events (Fig. 4D; hCohesin + Scc2/4) detected with yeast cohesin tetramers (Fig. 2C: 125mM). Therefore, hCohesin tetramers containing STAG1 have conserved the ability to bridge DNA. Importantly, hCohesin was able to form both “reversible” and “permanent” bridges (Fig. 4D; hCohesin + Scc2/4).

Besides mediating sister chromatid cohesion (*1, 2*), cohesin hold individual chromatids in *cis* forming loops (*4, 27, 28*). Recently, yeast condensin was the first SMC complex shown to exhibit an activity compatible with loop extrusion (*7*). It is unclear whether this activity is also present in the other eukaryotic SMC complexes; cohesin and Smc5/6. Condensin loop extrusion activity leads to compaction of linear DNA against forces of up to 2pN in magnetic tweezers (*17*). We purified yeast condensin (Supplementary Fig. 13) using an established protocol (*18, 29*) and tested whether it could compact λ-DNA molecules extended in the optical tweezers against a force of 1pN. A single λ-DNA molecule was first captured between the beads. We then immobilised one of the beads and applied a constant force of 5pN to the other bead in the opposite direction. This maintains to DNA extended with ∼14 µm between beads. We then moved the DNA to a channel containing 1nM condensin in 50 mM NaCl buffer supplemented with 1mM ATP and we incubated it for 30s. We then moved the DNA to a different channel containing 1mM ATP in 50mM NaCl buffer and reduced the extension force to 1pN. We recorded the distance between the two beads over time (Fig. 4E; condensin). We observed a progressive decrease of the distance between the beads (Fig. 4E; condensin, and Supplementary Fig. 14), consistent with the activity of condensin as a motor that compacts DNA (*17*). Some condensation events occurred in short bursts and caused the molecule to shorten ∼1-2µm in a few seconds (Fig. 4E; condensin, and Supplementary Fig. 14). The compaction rates of these events are compatible with the loop extrusion rates of 1500 base pairs per second reported recently (*7*), suggesting that the shortening of the molecule might indeed occur through this mechanism. After incubation, we generated a FE curve which showed the presence of sawtooth peaks characteristic of protein-mediated DNA bridging (Fig. 4E; bottom) (*23, 24*). Importantly condensin bridges were fully reversible and disappeared when the DNA was extended (Fig. 4E; bottom), consistent with the possibility that DNA loops had indeed formed through loop extrusion. Next, we sought to text whether yeast cohesin tetramers could also compact extended λ-DNA molecules in this assay. We incubated the DNA extended using 5pN of force with 1 nM cohesin, 2.5nM Scc2-Scc4 complex and 1mM ATP in 50mM NaCl buffer initially and moved the extended DNA to a buffer only channel (1mM ATP in 50mM NaCl) were the extension force was reduced to 1pN. We then recorded the distance between the two beads over time (Fig. 4F; cohesin). The distance between the beads did not change during the course of the experiment (Fig. 4F; cohesin and Supplementary Fig. 15), therefore we have to conclude that cohesin tetramers do not exhibit DNA compaction activity in this assay. As expected, the FE curve generated after incubation showed no evidence of protein-mediated DNA bridging (Fig. 4F; bottom).

## Discussion

Kimura *et al.* first proposed that the SMC complex condensin might generate DNA loops (*5*). This proposal was aimed at explaining the introduction of (+) writhe by condensin in circular plasmids (*5*) and was based on an earlier model of “loop expansion” that was proposed for bacterial MutS action (*30*). Interestingly, MutS loop expansion was shown to occur as a consequence of ATP-dependent bidirectional movement of the MutS dimer from the initial loading site (*30*). Although we did not detect DNA compaction by yeast cohesin tetramers, as predicted from a potential loop extrusion activity, we cannot rule out that cohesin extrudes DNA loops when additional factors are present. It is important to consider that HiC data demonstrates that removal of cohesin leads to loss of contacts at TAD boundaries (*6, 31, 32*), demonstrating that the complex is required for the maintenance of TAD signals. The loop extrusion activity for cohesin is a model conceived as an explanation for the enrichment in convergent orientation of CTCF motifs at TAD boundaries. However, nature has evolved alternative mechanisms that bias interactions of distant regions with preferred sequence orientations (*33*). Loop extrusion activity has been so far demonstrated for yeast condensin (*7*), further experiments will be required to directly test whether or not cohesin complexes have the ability to loop extrude.

In summary, here we have shown that cohesin complexes can form different types of bridges between dsDNAs and that this activity requires Scc2-Scc4 and ATP. Our results using two DNA molecules demonstrate that “permanent” cohesin tethers can move when force is applied (Fig. 4A), however, when the “permanent” bridges occur in *cis*, cohesin complexes cannot slide off the DNA molecules (Fig. 2C and Supplementary Fig. 8). The simplest explanation is that the two DNA molecules tethered are not located in the same physical space within the protein. The two main models proposed to explain how cohesin holds sister chromatids are the “ring” and “handcuff” models. The basic difference between these two models is the fact that in the ring model, the two DNAs occupy the same physical space within cohesin, i.e. they are co-entrapped in one compartment of the cohesin structure (*8, 9*), while in the “handcuff” model (and all its variations) the two DNAs are located in different physical compartments (*1, 10, 11*), generally argued to be two separate (but interacting) complexes. Based on the ring model it would be expected that cohesin slides off molecules when bridging them intramolecularly (Supplementary Fig. 15), however, our observations suggest that this is not the case (Fig. 2C and Supplementary Fig. 16). We thus propose that cohesin holds DNAs together by a handcuff-like mechanism (involving one or two cohesin complexes) where two physically separated compartments are involved in DNA tethering (Fig. 4G and Supplementary Fig. 16). This could be two compartments within one cohesin tetramer (Fig. 4G; 3-bar-buckle and “multiple subcompartments” models) or different compartments of two cohesin complexes (Fig. 4G; handcuff model). Similarly, in the single complex option the DNAs could be held in separated compartments (i.e. multiple subcompartments-pretzel-model), or could occupy two compartments jointly (i.e. 3-bar-buckle model). The activities described here are fully consistent with the original role attributed to cohesin in maintaining sister chromatid cohesion (*1, 2*). Therefore, our work provides a new critical tool for future investigations to further decipher how cohesin executes one of the critical functions required for genome inheritance, i.e. maintaining sister chromatids in close proximity from the time they are born in S phase until they are separated in anaphase.

## Acknowledgements

We would like to thank Jordi Cabanas Danés and Johanna Andrecka (LUMICKS) for technical help. We would like to thank our laboratory members for discussion and critical reading of the manuscript. We would like to thank D. D’Amours, C.Haering and J.Peters for sharing plasmids for the expression of yeast condensin and human cohesin. We thank Zdeněk Lánský for facilitating access to a quadrupole optical tweezer setup. The work in the L.A. laboratory was supported by Wellcome Trust Senior Investigator award to L.A. (100955, “Functional dissection of mitotic chromatin”) and the “London Institute of Medical Research (LMS), which receives its core funding from the UK Medical Research Council. J.S. acknowledges funding from the Deutsche Forschungsgemeinschaft (DFG) under grant STI673-2-1 and from the European Research Council under the ERC grant agreement 758124. The Single Molecule Imaging Group is funded by a core grant of the MRC-London Institute of Medical Sciences (UKRI MC-A658-5TY10), a Wellcome Trust Collaborative Grant (P67153), and a BBSRC CASE-studentship (to MDN).

## Author contributions

P. G-E. expressed and purified yeast cohesin and Scc2-Scc4 proteins and performed biochemical assays. J.D. expressed and purified yeast condensin. I. A. expressed and purified human cohesin. P. G-E., M.N and A.LL. collected optical tweezers datasets. P.G-E, M.N. and J.S. processed optical tweezers data. J. H. and L.T. performed ATPase assays. R. A prepared EM grids and collected and processed EM images. H.K. and A.M. performed mass spectrometry analysis. J. H. and J.S. performed, collected and analysed DNA curtain datasets. P. G-E. and L.A. conceived the project. L.A. wrote the manuscript. L.A., D.R and J.S revised the manuscript.

## SUPPLEMENTARY MATERIALS

Materials and Methods.

Supplementary Figures 1-16.

Table S1 and S2.

