## Supplementary materials and figures for "A conserved activity for cohesin in bridging DNA molecules"

**SUPPLEMENTARY INFORMATION**

**Methods**

**Protein expression and Purification**

The different subunits of the *S. cerevisiae* Scc2-Scc4 and cohesin complexes were synthesized under the control of galactose inducible promoters and cloned into multicopy episomal vectors (*URA3-SCC4-GAL1-10promoter-SCC2*-3xmyc-3xStrepII*;TRP1-SMC1*-3xStrepII-*GAL1-10promoter-SMC3 GAL7promoter-MCD1*-8xHis-3xHA; *URA3-GAL1-10promoter-SCC3*). Yeast W303-1a strains carrying the different constructs (CCG14800 for Scc2-Scc4 complex, CCG14801 for cohesin tetramer, and CCG14815 for cohesin *smc3*-K38I tetramer) were grown at 30°C in selective dropout media containing 2% raffinose and 0.1% Glucose to $OD_{600}$ of 1. Protein expression was induced by addition of 2% galactose and cells were grown for further 16 hours at 20°C. Cells were then harvested by centrifugation at 4°C, resuspended in two volumes of buffer A (25mM Hepes pH 7.5, 200 mM NaCl, 5% glycerol, 5 mM β-mercaptoethanol) containing 1× cOmplete EDTA-free protease-inhibitor mix (Roche), frozen in liquid nitrogen and lysed in a FreezerMill (SPEX Certiprep 6870). Cell powder was thawed at 4°C for 2 hours before mixing it with one volume of buffer A containing benzonase (Millipore) and incubated at 4°C for an extra hour. Cell lysates were clarified by centrifugation at 45 000 g for 1 hour followed by filtration using 0.22µm syringe filters. Clarified lysates were loaded onto 5ml StrepTrap-HP columns (GE Healthcare) pre-equilibrated with buffer A. The resin was washed with 5 column volumes of buffer A and eluted with buffer B (buffer A containing 5mM desthiobiotin). The peak fractions containing the over-expressed proteins were pooled together and salt concentration was adjusted to 150mM NaCl using 100mM NaCl-buffer A. Samples were then filtered as described above to remove residual aggregates and loaded onto 5ml HiTrap Heparin HP (GE Healthcare) columns pre-equilibrated with 150mM NaCl-buffer A. Elution was carried out using a linear gradient from 150 mM to 1 M NaCl in buffer A. Peak fractions were pooled and concentrated by centrifugal ultrafiltration (100kDa Amicon Ultra, Millipore). Salt concentration was adjusted to 300mM NaCl during the concentration step. Gel Filtration was carried out using a Superose 6 Increase 100/300 GL column (GE Healthcare) in 300mM NaCl buffer A. Fractions corresponding to monomeric complexes were pooled and concentrated as described above. Purified proteins were analyzed by SDS PAGE (NuPAGE 4-12% Bis-Tris protein gels, ThermoFisher Scientific) and Coomassie staining (InstantBlue, Expedeon). Protein identification was carried out by mass spectrometry analysis and Western blot. *S. cerevisiae* condensin complex was expressed and purified as previously described (*1, 2*)

Human cohesin tetramer was purified as described before (*3*). Human cohesin subunits (SCC1, 10xHis-SMC1A, SMC3-FLAG, SA1) were co-expressed in High Five insect (BTI-Tn-5B1-4) cells. Cells were disrupted by short sonication. Afterwards, the lysate was clarified by high-speed centrifugation. The complex was then purified via HisTrap (washing buffer: 25mM Tris pH 7.5, 500mM NaCl, 5% Glycerol, 2mM MgCl2, 20mM Imidazole, 0.01% Tween 20, 20mM ß-ME; elution buffer: 25mM Tris pH7.5, 150mM NaCl, 5% Glycerol, 2mM $MgCl_2$, 150 mM Imidazole, 0.01% Tween 20). Fractions were pooled and dialysed (25mM Tris pH 7.5 150mM NaCl 5% Gly 2mM $MgCl_2$). The protein was further purified by tandem ion exchange chromatography using first an anion exchange column connected to a cation exchange column. The complex was then eluted from the cation exchange column (25mM Tris pH 7.5, 1M NaCl, 5% Glycerol, 2mM MgCl2). Subsequently, the peak fractions were pooled and dialyzed into storage buffer (25mM Tris pH 7.5, 150mM NaCl, 5% Glycerol, 2mM $MgCl_2$). Purity was confirmed by gel electrophoresis and mass spectrometry.

**Electrophoretic gel mobility shift assay**

Increasing concentrations of Scc2–Scc4 complex ranging from 100 to 800nM were incubated for 45 min with 50ng of pUC19 at 30°C in 25 mM Tris–HCl pH 7.0, 50 mM NaCl, 8% glycerol, 0.1 mg/ml BSA and 0.5 mM DTT in a final volume of 15µl. The reactions were resolved by electrophoresis for 1 h at 80V on 0.8% (w/v) TAE-agarose gels at 4°C. DNA was detected on a fluorescent image analyzer FLA-5000 (Fujifilm) after SYBR Green I (Invitrogen, ThermoFisher Scientific) gel staining. Condensin assays were carried out as previously described (*2*).

**Protein cross-linking and Electron Microscopy**

For cross-linking of cohesin complex, protein samples were incubated with BS3 at a 1:3000 molar ratio in buffer XL (25 mM Hepes, 125 mM NaCl, 5% glycerol, 1mM DTT, pH 8) for 2 hours on ice before quenching with 10 mM Tris-HCl pH8 for 30 min on ice.

Negative stain grids were prepared as follows: 3.5 µl of suspended sample (final concentration of 0.02 mg/ml in buffer XL) were deposited on glow-discharged grids coated with a continuous carbon film. The sample was left on the grid for one minute before blotting the excess liquid. A 3.5 µl drop of 2% uranyl acetate solution was added for 1 minute, the stain was blotted away and the grids left to dry.

A set of 250 micrographs was collected on a Philips CM200 TWIN FEG electron microscope operated at 160 kV (MRC LMS cryo-EM facility). Images were recorded on a Tietz 2k CCD camera at a nominal magnification of 38,000 and a final pixel size of 3.58 Å. CTF parameters were estimated using Gctf[2]. A total of ~9,000 particles were automatically picked using Gautomatch software (K. Zhang, unpublished) using class averages obtained from a manually picked subset of 1,500 particles as references. The following 2D classifications were performed with RELION v3.0 beta(*5*).

***In vitro* cohesin loading assay**

Cohesin loading assays were done as described in (*6*) using pUC19 plasmid. Topologically bound DNA-Cohesin complexes were immunoprecipitated using a µMACS HA isolation kit (Miltenyi Biotec). Following incubation with PstI and/or protein digestion, the recovered DNA was analyzed by electrophoresis on a 0.8% (w/v) TAE-agarose gel in 1xTAE and visualized as described above.

**ATPase assays**

For the ATPase assays, 30nM cohesin was incubated at 29°C with 60nM Scc2/4 and 0.2nM λ-DNA (NEB) in ATPase Buffer (35 mM Tris-HCl pH 7.0, 20 mM NaCl, 0.5 mM $MgCl_2$, 13.3 % Glycerol, 0.003 % Tween 20, 1 mM TCEP, 0.2 mg/ml BSA). The reaction was started by adding 400 µM ATP spiked with [γ-32P]-ATP. 1 µl samples were taken after 1, 15, 30 and 60 min. The reaction was immediately stopped by adding 1µl 50mM EDTA before spotting the samples on polyethylenimine-cellulose F sheets. The free phosphate was separated from ATP using thin layer chromatography with 0.5M LiCl, 1M formic acid as the mobile phase. The spots were detected on a phosphor imager and analyzed using ImageJ. Data points were corrected for spontaneous ATP hydrolysis. Each reaction was performed in triplicate. Data were fitted to Michaelis-Menten kinetics.

**Single molecule experiments**

DNA Curtains experiments were performed as described previously (*7*). Briefly, flow cells were produced by deposition of chromium features onto fused silica microscope slides by e-beam lithography. Flow cells were connected to a microfluidics system based on a syringe pump (Landgraf GmbH) and two injection valves (Idex) and illuminated by 488nm or 561nm lasers (Coherent) in a prism-type TIRF configuration on an inverted microscope (Nikon Ti2e). Imaging was performed by an EMCCD camera (Andor iXon life) with illumination times of 100ms. λ-DNA (NEB) was end-modified by hybridizing biotinylated or digoxigeninated oligos complementary to the *cos* site and purified by size exclusion chromatography. Modified lambda DNA was anchored to the surface of a lipid bilayer in flow cells by biotin-streptavidin-biotin interactions, stretched by flow across chromium barriers and anchored to downstream chromium pedestals by the digoxigenin-binding protein DIG10.3 (*8*). Experiments were performed in buffer M (40mM Tris-HCl pH 7.8, 1mM MgCl2, 1mM DTT, 1mg/ml BSA, 0.16nM YOYO-1). Cohesin complexes were labeled by incubating them at a concentration of 3nM in a small volume of buffer M supplemented with 50mM NaCl with 3x molar excess Qdots (SiteClick 705 kit, Invitrogen) fused to anti-HA antibodies (3F10, Roche) for 30 minutes at 4C. The mixture was then supplemented with 8nM Scc2/Scc4, 100$\mu$m biotin and 0.5mM of nucleotide (ATP, ADP or ATP$\gamma$S), if required, prior to injection. For diffusion measurements, the flow cell was flushed after the completion of loading with buffer M supplemented with KCl at the indicated concentrations and the flow was stopped. Illuminations were performed either continuously (diffusion and lifetime measurements) or with lower frame rates (inter-molecular bridging videos). To minimize photo-damage, 488nm pulses to illuminate the DNA, if required, were only used at every 10$^{th}$ illumination.

Videos were recorded in NIS Elements (Nikon) and analysed using custom-written software in Igor Pro (Wavemetrics). Lifetime measurements and initial binding distributions of cohesin complexes on DNA were generated by manually analysing kymograms. Survival curves were generated by a Kaplan-Meier estimator, bootstrapped, and fitted to a double-exponential model.

For the determination of diffusion coefficients, labelled cohesin complexes were tracked using custom-written software and the diffusion coefficients were extracted using a maximum-likelihood estimator (*9*) as described previously (*10*).

Optical tweezers experiments were carried out on C-trap and Q-trap systems integrating optical tweezers, confocal fluorescence microscopy and microfluidics, and recorded using BlueLake software (LUMICKS). The laminar flow cell was passivated using 0.50% Pluronic and 2mg/ml BSA. Biotin-labelled double-stranded λ-DNA molecules were tethered between two streptavidin coated, polyesterene beads (4.42 $\mu$m in diameter, Spherotech). Depending on the experiment, one or two individual double-stranderd λ-DNA molecules were attached between two beads. The beads were previously passivated with 1mg/ml BSA. After DNA capture, beads were incubated inside the protein channel either in a relaxed (~3$\mu$m apart) or extended position (Force Clamp at 5pN, ~14$\mu$m apart) for 30 seconds and then returned to the buffer channel for force-extension (FE), force clamp, and fluorescence analysis. Cohesin and Scc2-Scc4 complex were used at 1nM and 2.5nM concentration, respectively. Beads and DNA catching, and protein loading were performed in a buffer containing 50 mM Tris-HCl pH 7.5, 50 mM NaCl, 2.5mM MgCl2, 0.5mg/ml BSA, 40 $\mu$M biotin and 1mM DTT. When indicated, ADP, ATP-$\gamma$-S or ATP were added to both protein and buffer channels at a final concentration of 1mM. Salt concentration was modified from 50mM to 125, 300 or 500mM NaCl in the buffer channel as specified in the text and figures. FE curves were performed at a speed of 1$\mu$m/s. Compaction experiments were carried out at a constant force of 1pN. For friction experiments, beads were moved 6$\mu$m, back and forth, at a speed of 0.2$\mu$m/s. SYTOX Orange (Invitrogen, ThermoFisher Scientific) was used at a final concentration of 50mM for DNA imaging, using a 532nm wavelength laser. Force data was processed using Igor Pro 7 software (Wavemetrics) and images using Adobe Photoshop CC.

**Western Blot**

For Western blot, 2 µg of purified complexes were run on NuPAGE 4-12% Bis-Tris gels (ThermoFisher Scientific), transferred to Immobilon-P membranes (Millipore) and probed with anti-Strep (ab180957, Abcam, 1:5000) and anti-HA (3F10, Roche, 1:5000) antibodies in 5%milk-PBS 0.01% Tween overnight at 4ºC. Membranes were then washed and incubated with HRP anti-rabbit (Santa Cruz Biotechnology, 1:40 000) and anti-rat (Jackson ImmunoResearch, 1:10 000) secondary antibodies respectively for 1 hour at room temperature. Immunoblots were developed using the Luminata Forte detection reagent (Millipore) and Hyperfilms ECL (GE Healthcare).

**Liquid chromatography-tandem mass spectrometry (LC-MS/MS)**

Samples were processed by in-Stage Tip (iST) digestion (Preomics GmbH, Planegg/Martinsried) following the manufacturer recommendation. Protein digests were solubilised in 30 µl of reconstitution buffer and were transferred to auto sampler vials for LC-MS analysis. Peptides were separated using an Ultimate 3000 RSLC nano liquid chromatography system (Thermo Scientific) coupled to a LTQ Orbitrap Velos mass spectrometer (Thermo Scientific) via an EASY-Spray source. Sample volumes were loaded onto a trap column (Acclaim PepMap 100 C18, 100 µm x 2 cm) at 8 µl/min in 2% acetonitrile, 0.1% TFA. Peptides were eluted on-line to an analytical column (EASY-Spray PepMap C18, 75 µm x 50 cm). Peptides were separated using a ramped 120 min gradient from 1-42% buffer B (buffer A: 5% DMSO, 0.1% formic acid; buffer B: 75% acetonitrile, 0.1% formic acid, 5% DMSO). Eluted peptides were analysed operating in positive polarity using a data-dependent acquisition mode. Ions for fragmentation were determined from an initial MS1 survey scan at 30,000 resolution (at m/z 200) in the Orbitrap followed by CID (Collision-Induced Dissociation) of the top 10 most abundant ions in the

Ion Trap. MS1 and MS2 scan AGC targets set to 1e6 and 1e5 for a maximum injection time of 50 ms and 110 ms, respectively. A survey scan m/z range of 350 – 1500 m/z was used, with CID parameters of isolation width 1.0 m/z, normalised collision energy of 35%, activation Q 0.25 and activation time of 10ms.

Data were processed using the MaxQuant software platform (v1.6.2.3) with database searches carried out by the in-built Andromeda search engine against the Uniprot Saccharomyces cerevisiae database (6,729 entries, v.20180305). A reverse decoy database was created and results displayed at a 1% false-discovery rate (FDR) for peptide spectrum matches and protein identification. Search parameters included: trypsin, two missed cleavages, fixed modification of cysteine carbamidomethylation and variable modifications of methionine oxidation, asparagine deamidation and protein N-terminal acetylation. Label-free quantification was enabled with an LFQ minimum ratio count of 2. 'Match between runs' function was used with match and alignment time limits of 0.7 and 20 min, respectively. Protein and peptide identification and relative quantification outputs from MaxQuant were further processed in Microsoft Excel, with hits to the 'reverse database', 'potential contaminants' (peptide list only) and 'Only identified by site' fields removed.

**Supplementary Figure Legends**

**Supplementary Figure 1. Purification of budding yeast Cohesin. A**. Purified cohesin tetramer containing Smc1, Smc3, Mcd1 and Scc3 was analysed by SDS-PAGE electrophoresis followed by Coomassie Blue staining. Western analysis showing Smc1-strep and Mcd1 HA is included. **B**. Representative micrograph of a BS3-crosslinked cohesin sample observed in negative stain. Scale bar 50 nm. Right panel: class averages obtained with RELION. A set of the best ~5.000 particles were used for this classification. The size of the circular mask is 450 Å.

**Supplementary Figure 2. Purification of budding yeast Scc2-Scc4 complex. A.** Coomassie Blue staining of purified Scc2-Scc4 complex. Western analysis showing Scc2-strep is shown.
**B**. Electrophoretic mobility shift assays using pUC19 as substrate and the indicated Scc2-Scc4 complex concentrations.

**Supplementary Figure 3. Topological loading of yeast cohesin on plasmid DNA. A.** Agarose gel electrophoresis showing recovered DNA after cohesin loading and immunoprecipitation with and without 0.5mM ATP both in the presence and absence of Scc2/4 complex. Topological assays were done as in (6). **B**. Gel image of recovered DNA in supernatant (S) and cohesin-bound bead (B) fractions after linearisation of immunoprecipitated cohesin-bound DNA by PstI digestion.

**Supplementary Figure 4.** ATP hydrolysis by cohesin and cohesin ATPse mutant K38I with or without Scc2-Scc4 complex.

**Supplementary Figure 5. Analysis of yeast cohesin on DNA curtains.**
**A**. Image of cohesin tagged with quantum dots (QD) (magenta) bound at 25 mM KCl to λ-DNA before injection of high ionic strength buffer (top). Image of cohesin (magenta) bound to λ-DNA (green) after injection of 250 mM KCl buffer (bottom). **A**. Representative kymograph illustrating cohesin diffusion at 250 mM KCl. **C**. Distribution of bound cohesins on λ-DNA. A/T content is indicated. Error bars: 68% confidence intervals. Distribution of bound cohesins on flipped λ-DNA is shown (right graph). **D**.Correlation between cohesin localisation and A/T nucleotide content on λ-DNA (Pearson's r = 0.90). **E**. Initial binding positions of individual cohesin to λ-DNA. **F**. Diffusion coefficients for cohesin movement in buffers of different of ionic concentration.

**Supplementary Figure 6. Intramolecular cohesin bridging requires ATP.**
**A**. FE curves for λ-DNA pre-incubated with 1 nM cohesin, 2.5 nM Scc2-Scc4 complex and ATP analogues. After capturing a single DNA molecule between two optically trapped beads, DNA was incubated in the presence of protein in a relaxed conformation (3 μm bead distance) for 30s in 50mM NaCl and then moved to a buffer channel with 50mM NaCl for extension and measurements. Only ATP exhibits DNA bridging rupture events. Note that one of the FE curves (first curve in the ATP set)

was used to illustrate bond-rupture events in Fig. 2A. **B**. Distribution of cohesin-induced DNA bridge loop sizes calculated from bond-rupture events in FE curves.

**Supplementary Figure 7. Purification of budding yeast Cohesin ATPase mutant.** Purified cohesin tetramer containing Smc1, Smc3-K38I, Mcd1 and Scc3 was analysed by SDS-PAGE electrophoresis followed by Coomassie Blue staining. Western analysis showing Smc1-strep and Mcd1 HA is included.

**Supplementary Figure 8. Permanent cohesin bridges are not displaced by physical stretching of λ-DNA.** FE curves during sequential extension and relaxation cycles at 300mM and 500mM NaCl. After capture of λ-DNA between the two optically trapped beads, DNA is relaxed (3μm bead distance) and incubated for 30 seconds in the protein channel (1 nM cohesin and 2.5 nM complex and 1 mM ATP in 50mM NaCl). DNA is moved to a buffer channel (either 300mM NaCl or 500mM NaCl as indicated) before re-extension to test for DNA bridges. After confirmation of the bridges and full extension of the molecule (FE curves on the left), the DNAs are relaxed in the same channel (either 300mM NaCl or 500mM NaCl as indicated) and re-extended for a second time. FE curves of the second re-extension (FE curves on the right) show that the DNA bridge has not been displaced. Two independent molecules are shown. The first extension is also shown in Fig. 2D.

**Supplementary Figure 9. Cohesin does not capture two λ-DNAs in sequential steps.** λ-DNA incubated with 1nM cohesin, 2.5 nM Scc2-Scc4 complex and 1mM ATP in an extended conformation. The DNAs were moved to a buffer channel (50mM NaCl) in the presence of 1mM ATP (buffer only - dark blue- top two left FE curves) or 2.5nM Scc2-Scc4 complex and 1mM ATP (+Scc2/4 - light blue – bottom two left FE curves). The λ-DNA molecules were then incubated in a relaxed position (3μm bead distance) for 30s. DNAs were then moved to a only buffer channel (125mM NaCl containing 1mM ATP) and re-extended (right FE curves; top - only buffer - during the first relaxation, bottom buffer plus Scc2/4 during the first relaxation). The same molecules were then relaxed (3μm bead distance) and incubated (for 30s) with 1nM cohesin, 2.5 nM Scc2-Scc4 complex and 1mM ATP in 50mM NaCl. Finally, the DNA was moved to a different channel with 1mM ATP in 125mM NaCl and re-extended. FE curves of the final re-extension are shown (left FE curves; top- only buffer - during the first relaxation, -bottom- buffer plus Scc2/4 during the first relaxation). Only FE curves incubated with cohesin in a relaxed conformation show the presence of DNA bridging rupture events (i.e. FE curves on the right).

**Supplementary Figure 10. DNA friction experiments confirm the presence of cohesin complexes on extended λ-DNA.** Friction experiments were performed on a quadruple-trap optical tweezer system as described in (*11*). Two molecules of λ-DNA were tethered independently between two pairs of beads. Beads were then moved to the protein channel, containing 1nM cohesin and 2.5nM Scc2/4 complex in 50mM NaCl plus 1mM ATP, and incubated separated in an extended position (~14 μm bead distance). After incubation, the beads were moved to the buffer channel containing 125mM NaCl and 1mM ATP and crossed, so contact between both DNA molecules was established. Beads 1 and 2 were then moved simultaneously in the x axis to cause the sliding of one of the DNA molecules on top of the other. Forces in both beads 2 (sliding DNA molecule) and 3 (static DNA molecule) were recorded. A negative control with naked DNA is included (left panel). The presence of bound protein was identified by the appearance of abrupt changes in both force 2 and force 3 caused by the movement of beads 1 and 2 (right panel). For graphical representation, force data were downsampled to 100Hz.

**Supplementary Figure 11. Generation of permanent cohesin bridges using a quadrupole trap optical tweezer. A.** Schematic representation of the experimental design for the quadrupole trap optical tweezer to generate permanent cohesin bridges. First, a pair of λ-DNAs (DNA1 and DNA2) are trapped between two pairs of beads (beads 1 and 2 trap DNA1, while beads 3 and 4 trap DNA2) and kept extended (15μm bead distance). DNA2 is manipulated using beads 3 and 4, moved over DNA1 and positioned at a 90 angle, then relaxed (3μm bead distance). The crossed DNAs are moved to a channel containing 1 nM cohesin and 2.5 nM Scc2-Scc4 complex and 1mM ATP in 50mM NaCl and incubated for 30s. The crossed DNAs are moved to a channel containing 1mM ATP in 300mM NaCl and DNA2 is extended and moved back to its original position using beads 3 and 4. We used 50nM of SYTOX Orange to visualise the bridged DNA (right image). DNA molecules are in green, beads are numbered.

**Supplementary Figure 12. Purification of human Cohesin. A**. Purified cohesin tetramer containing hSmc1A, Smc3-FLAG, Scc1 and 10xHis SA1 was analysed by SDS-PAGE electrophoresis followed by Coomassie Blue staining. Purification was done using biGBac vector (pBIG1c) as described previously (*3*).

**Supplementary Figure 13. Purification of yeast condensin. A.** Purified condensin pentamers containing Smc2, Smc4, Brn1 and Ycs4 and Ycg1 was analysed by SDS-PAGE electrophoresis followed by Coomassie Blue staining. Purifications were done as in (*1, 2*). **B.** Electrophoretic mobility shift assays with a 6-carboxyfluorescein–labelled 41-bp dsDNA substrate (100 nM) and the indicated protein concentrations.

**Supplementary Figure 14. Budding yeast condensin but not cohesin compacts λ-DNA against 1pN stretching force.** Examples of DNA compaction traces for λ-DNA molecule extended using a force of 1pN (top). The DNAs were tethered between two beads. One bead was clamped (fixed) while a 5pN force was applied to the second bead to maintain the molecule extended. The DNA was then incubated in the presence of 1nM condensin (1mM ATP in 50mM NaCl) (top- condensin - magenta traces) or 1nM Cohesin and 2.5 nM Scc2-Scc4 complex (1mM ATP in 50mM NaCl) (bottom -cohesin -- yellow traces). Extended DNAs were then moved to a different channel containing 1mM ATP in 50mM NaCl and the extension force was reduced to 1pN. The distance between the beads was recorded over time. Only condensin was able to reduce the distance between the beads over time consistent with a DNA compaction activity. Two independent molecule traces for each complex is shown. An additional trace is shown in Fig. 4F-H. For graphical representation, force data were downsampled to 100Hz.

**Supplementary Figure 15. Expected behaviour of cohesin bridges from “ring” or “embrace” model. A.** Schematic representation of expected behaviour of intramolecular cohesin tethers from the proposed “ring” (or ”embrace”) model. The model proposes that cohesin co-entraps two DNAs within the its ring structure, i.e. both DNAs occupy one physical space within cohesin. From this model, it is expected that cohesin should be fully displaced from λ-DNA molecules when tethering in *cis* as force is applied to separate the beads (shown in the diagrammes). This is not what it is observed experimentally (Fig. 2C and Supplementary Fig. 8). **B.** Schematic representation of expected behaviour of intermolecular cohesin tethers from the proposed “ring” (or ”embrace”) model. The expectation from this model is that intermolecular cohesin bridges are able to move (slide) as illustrated in the diagramme. This is what it was observed experimentally (Fig. 4A).

**Supplementary Figure 16. Expected behaviour of cohesin bridges from “handcuff” or “pretzel” models. A.** Schematic representation of expected behaviour of intramolecular cohesin tethers from the “handcuff” and “multiple subcompartment” (“pretzel”) models. The “handcuff” model is based on the assumption that DNAs are located in different physical compartments, generally two distinct (but interacting) cohesin complexes. In the “multiple subcompartment” (“pretzel”) model, the DNAs are aloso located in different physical compartments, but within a single cohesin complex. The prediction from both models is that cohesin cannot be fully displaced from λ-DNA molecules when tethering them in *cis* (as shown in the illustration). This is what is observed experimentally (Fig. 2C and Supplementary Fig. 8). **B.** Schematic representation of expected behaviour of intermolecular cohesin tethers from the “handcuff” and “multiple subcompartment” (“pretzel”) model. The expectation from these models is that intermolecular cohesin bridges are able to move (slide) as illustrated in the diagramme. This is what it was observed experimentally (Fig. 4A).

**Supplementary Table 1.**
Protein identifications for cohesin tetramer (Wt and K38I) and Scc2-Scc4 purifications.

**Supplementary Table 2.**
Identifications of K38I peptides in purifications of cohesin ATPase mutant (Smc3-K38I).

#### Supplementary Figure 1

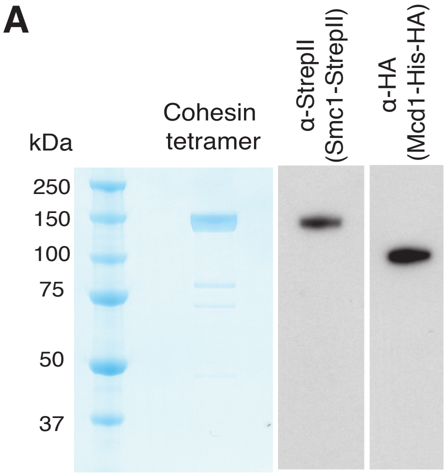


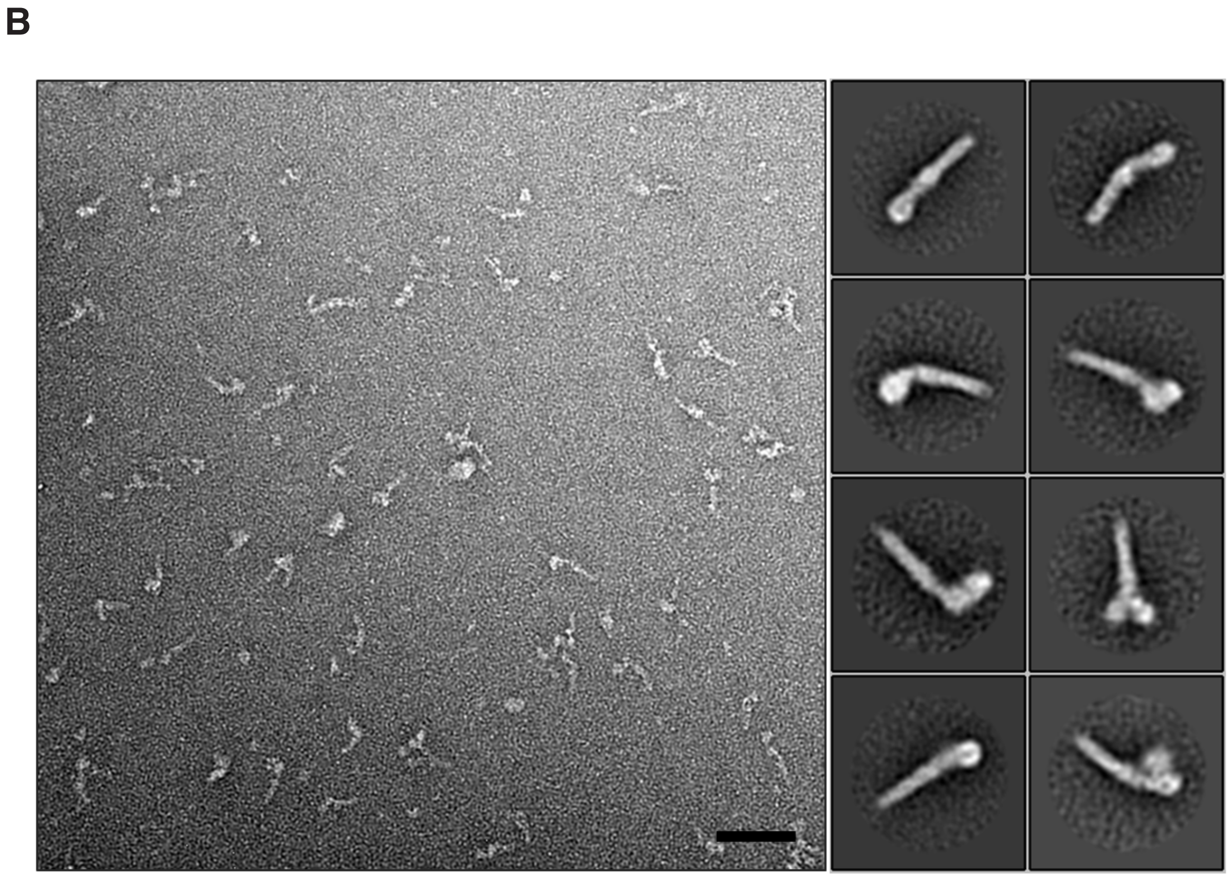

### Supplementary Figure 2

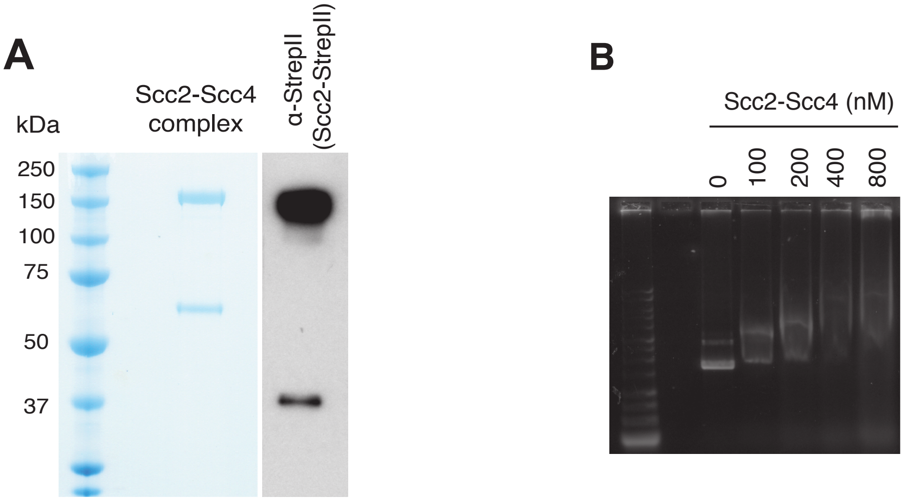

#### Supplementary Figure 3

A

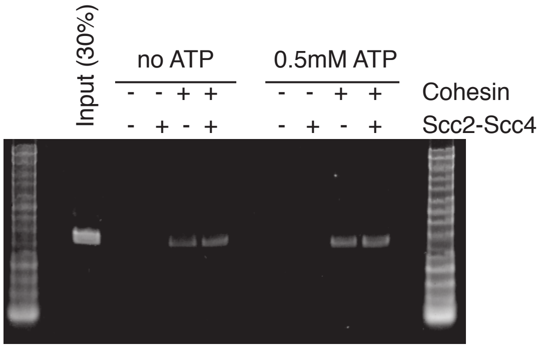


B

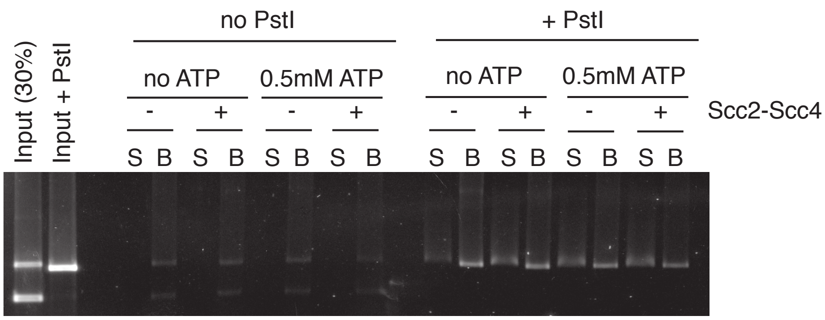

#### Supplementary Figure 4

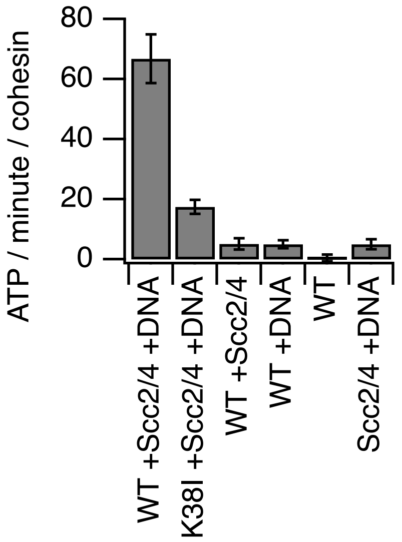


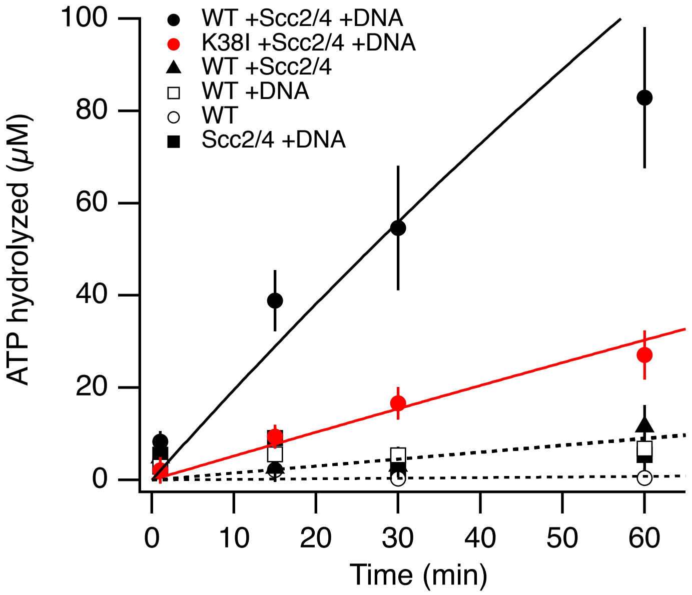

### Supplementary Figure 5

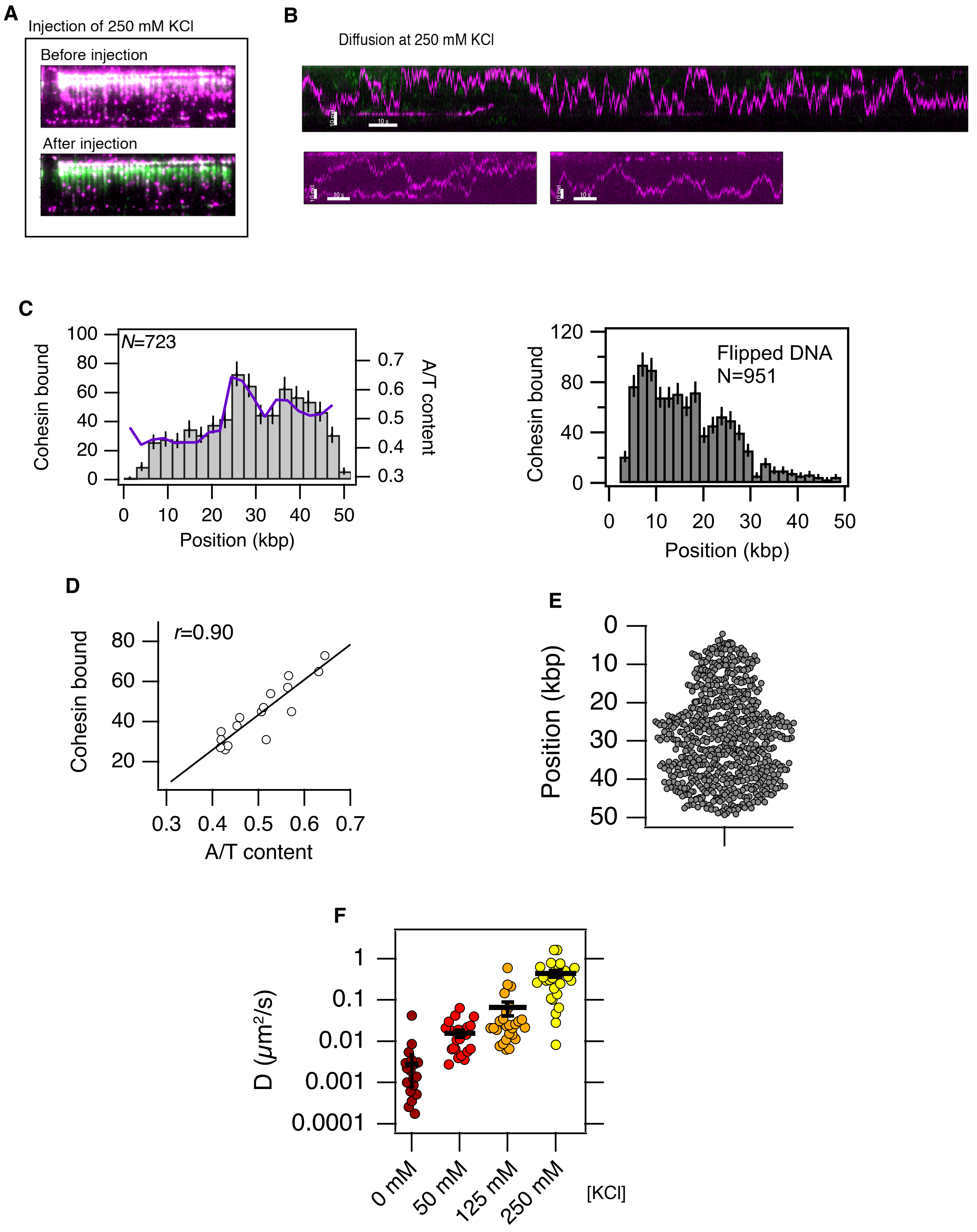

### Supplementary Figure 6

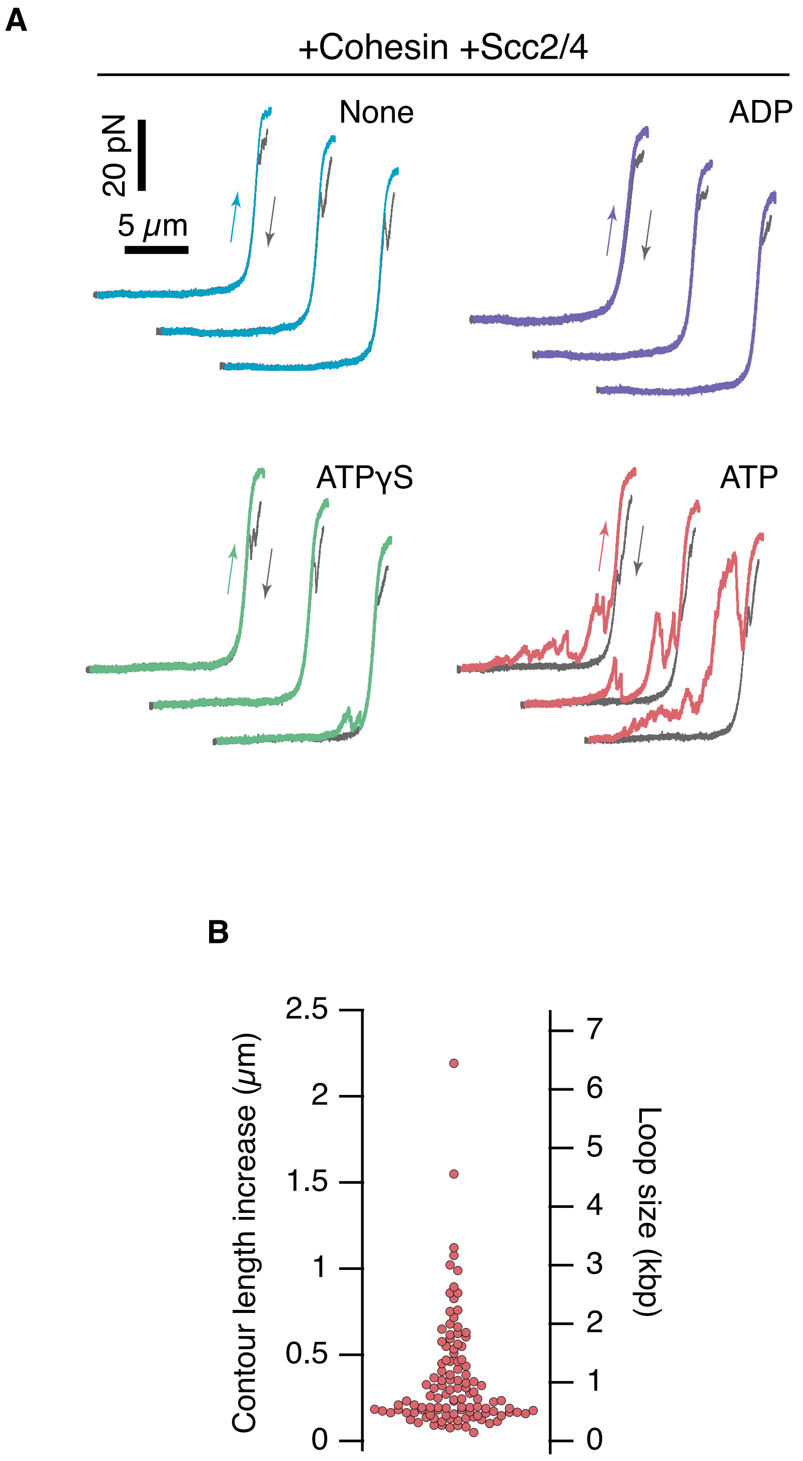

#### Supplementary Figure 7

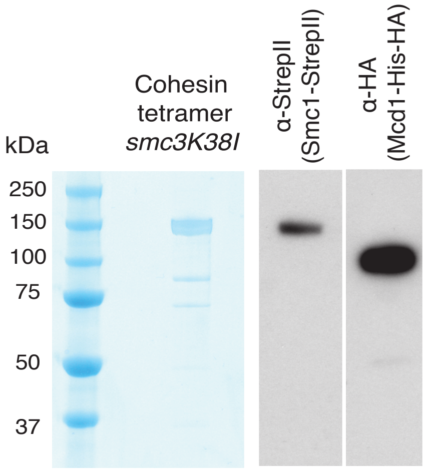

### Supplementary Figure 8

#### 300 mM NaCl

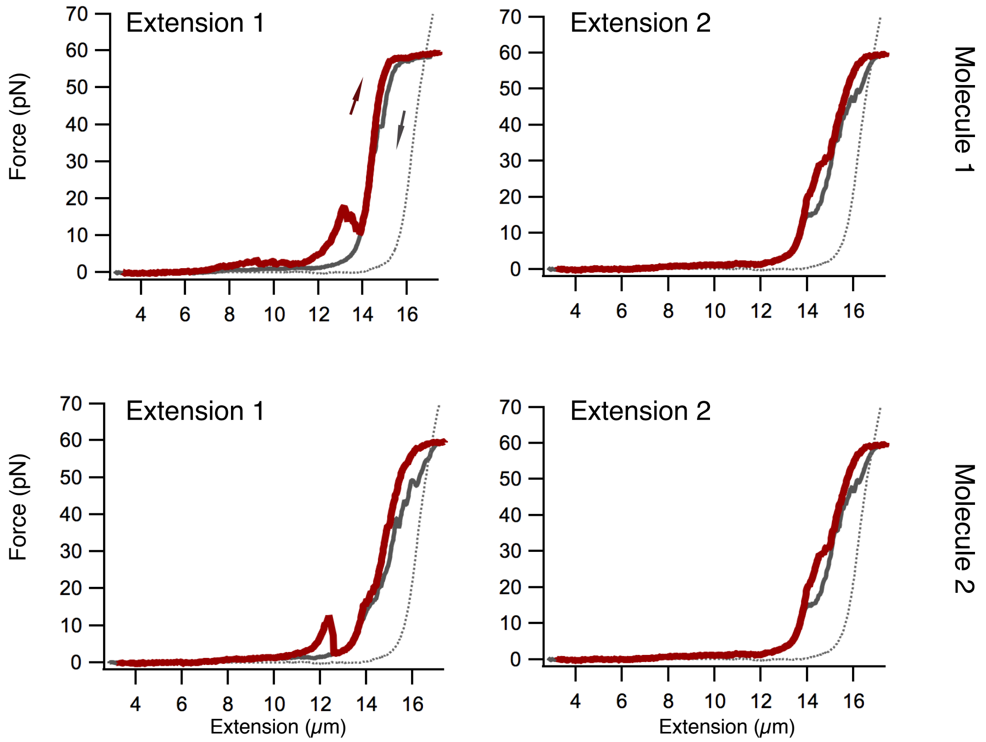


#### 500 mM NaCl

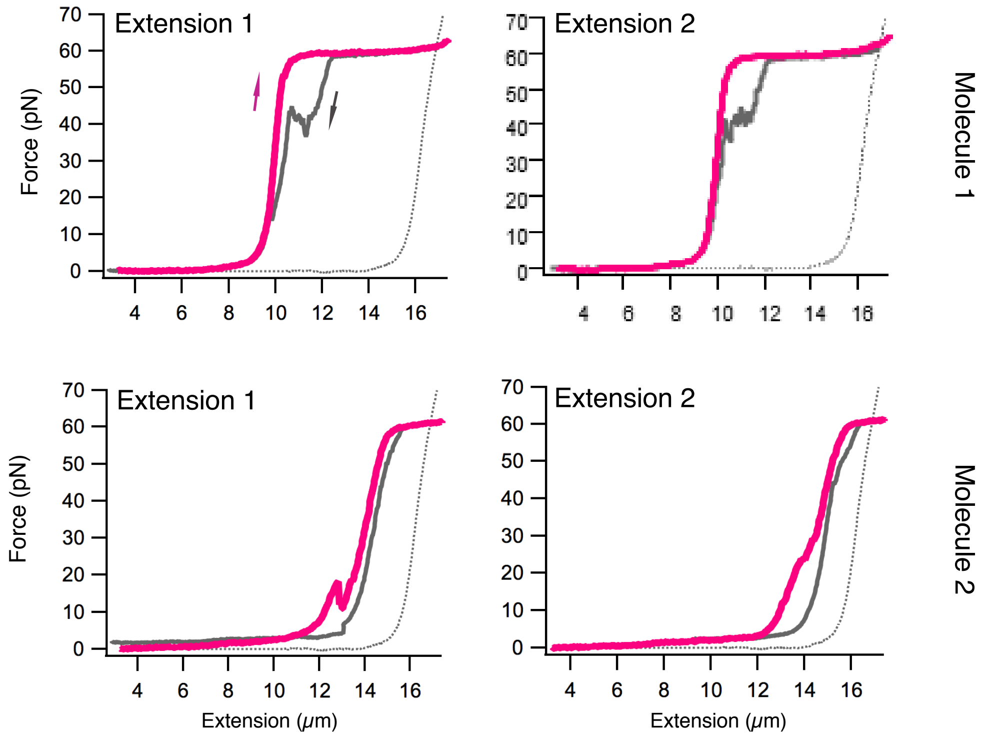

#### Supplementary Figure 9

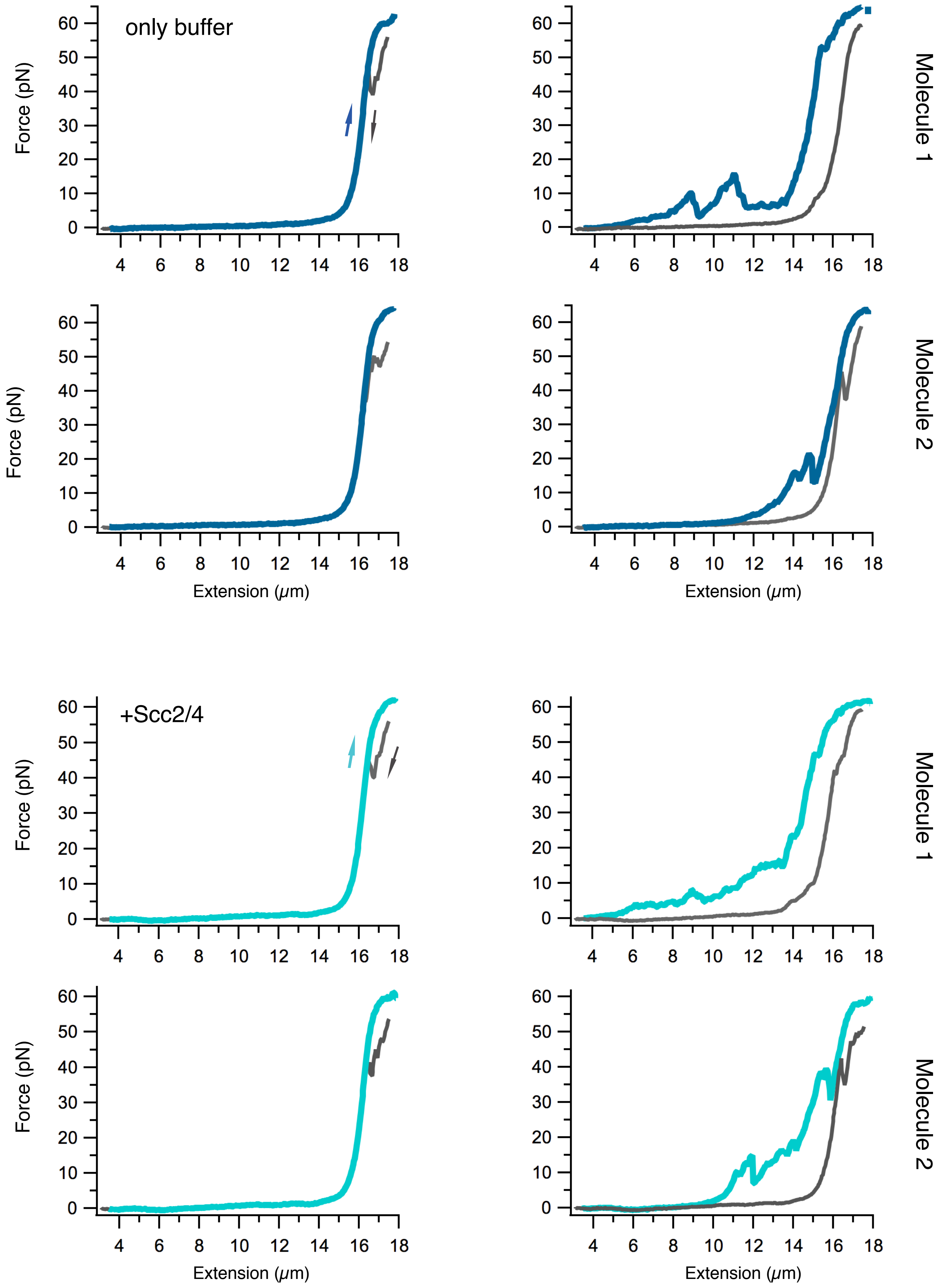

### Supplementary Figure 10

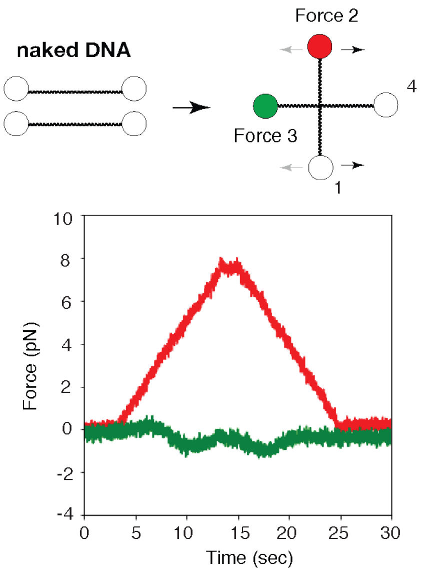


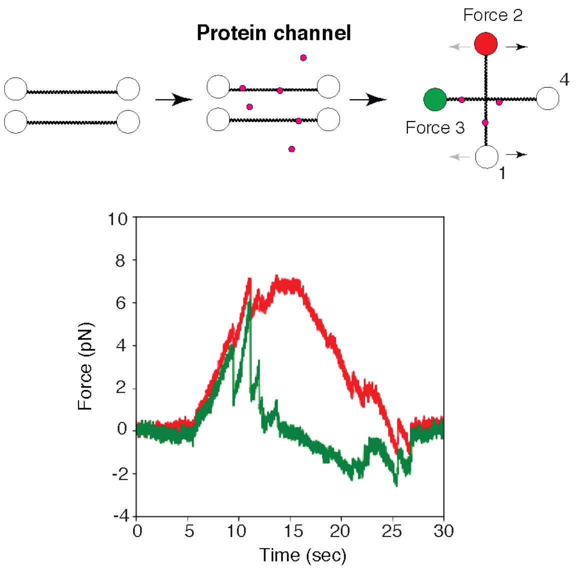

### Supplementary Figure 11

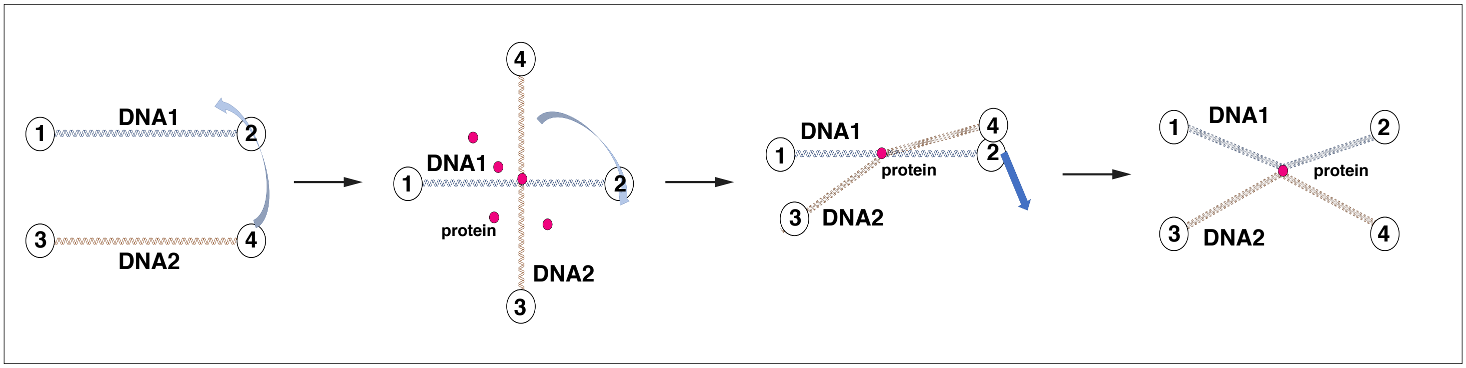


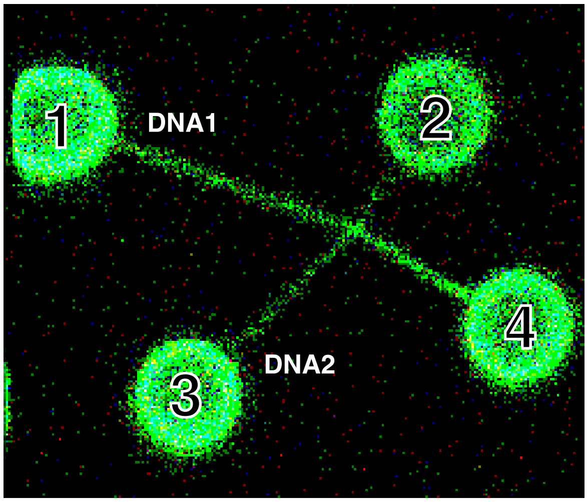

#### Supplementary Figure 12

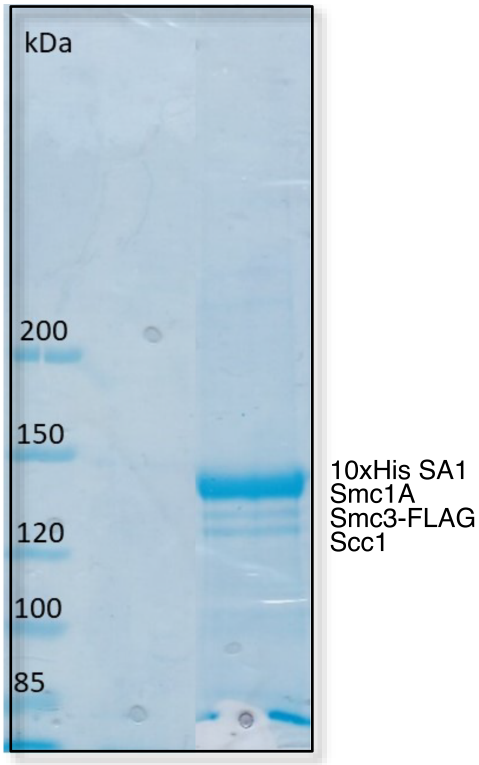

#### Supplementary Figure 13

A

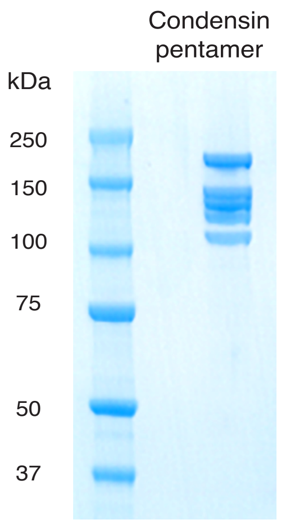


B

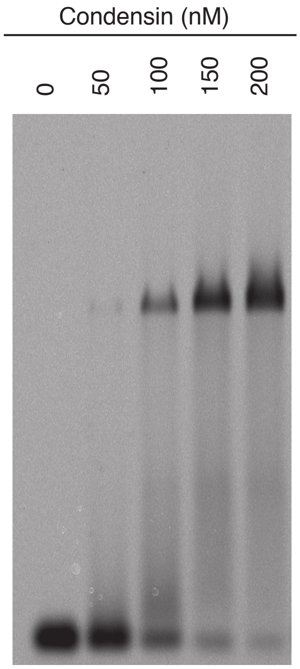

#### Supplementary Figure 14

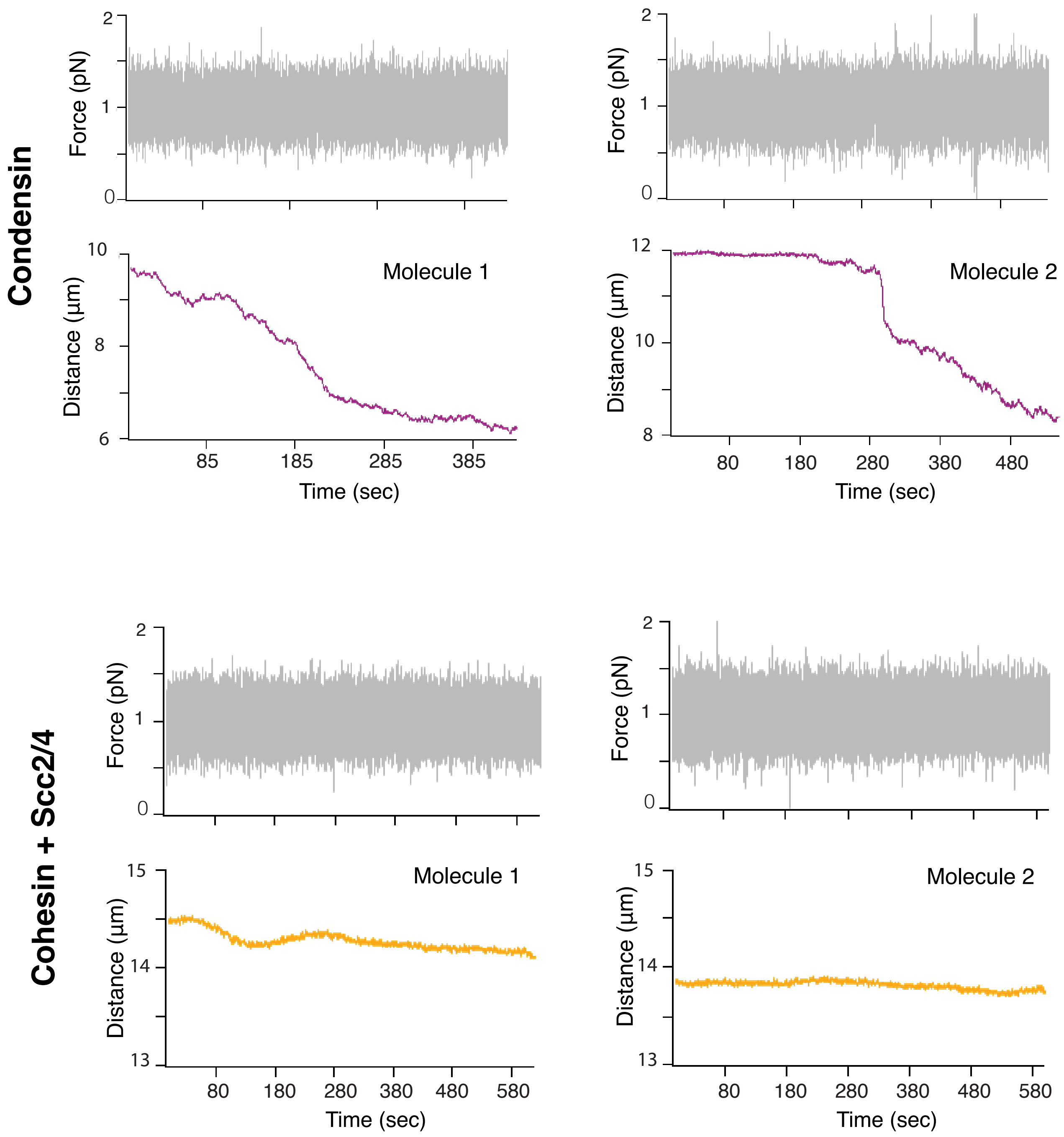

#### Supplementary Figure 15

**A**

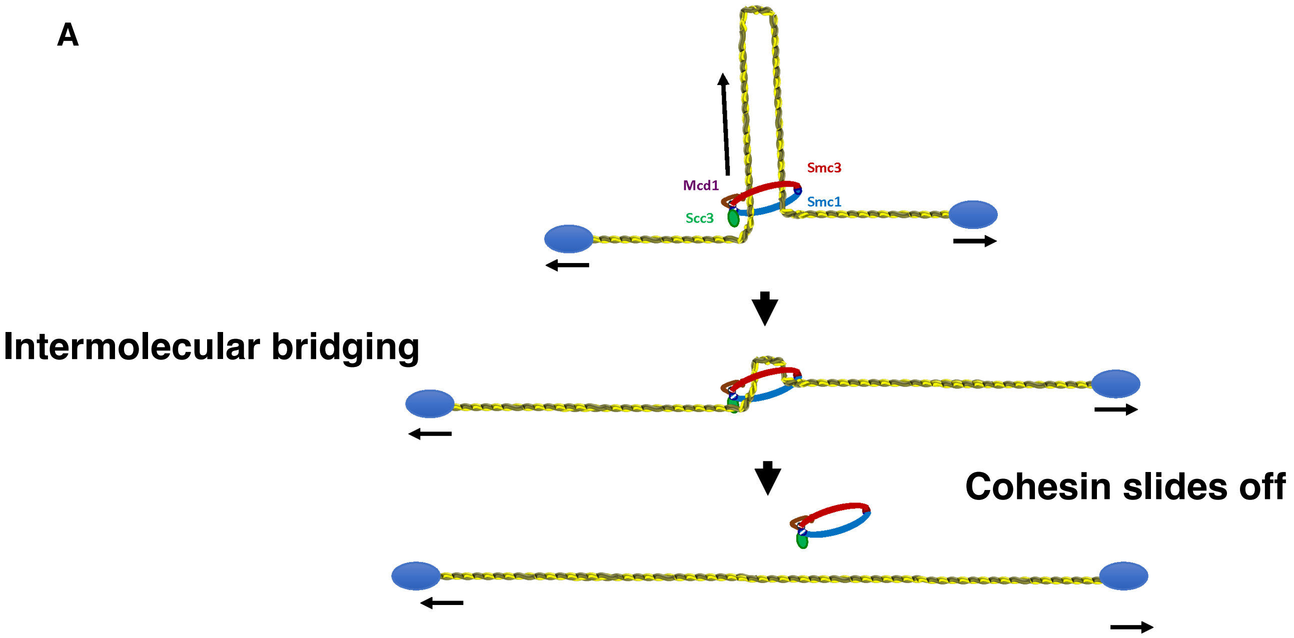


**B**

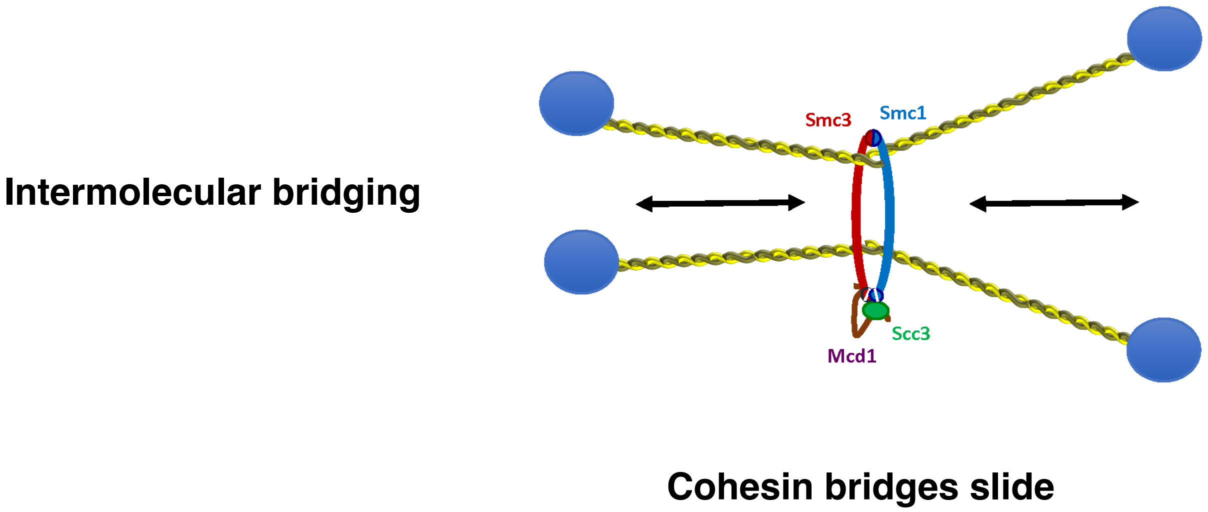

#### Supplementary Figure 16

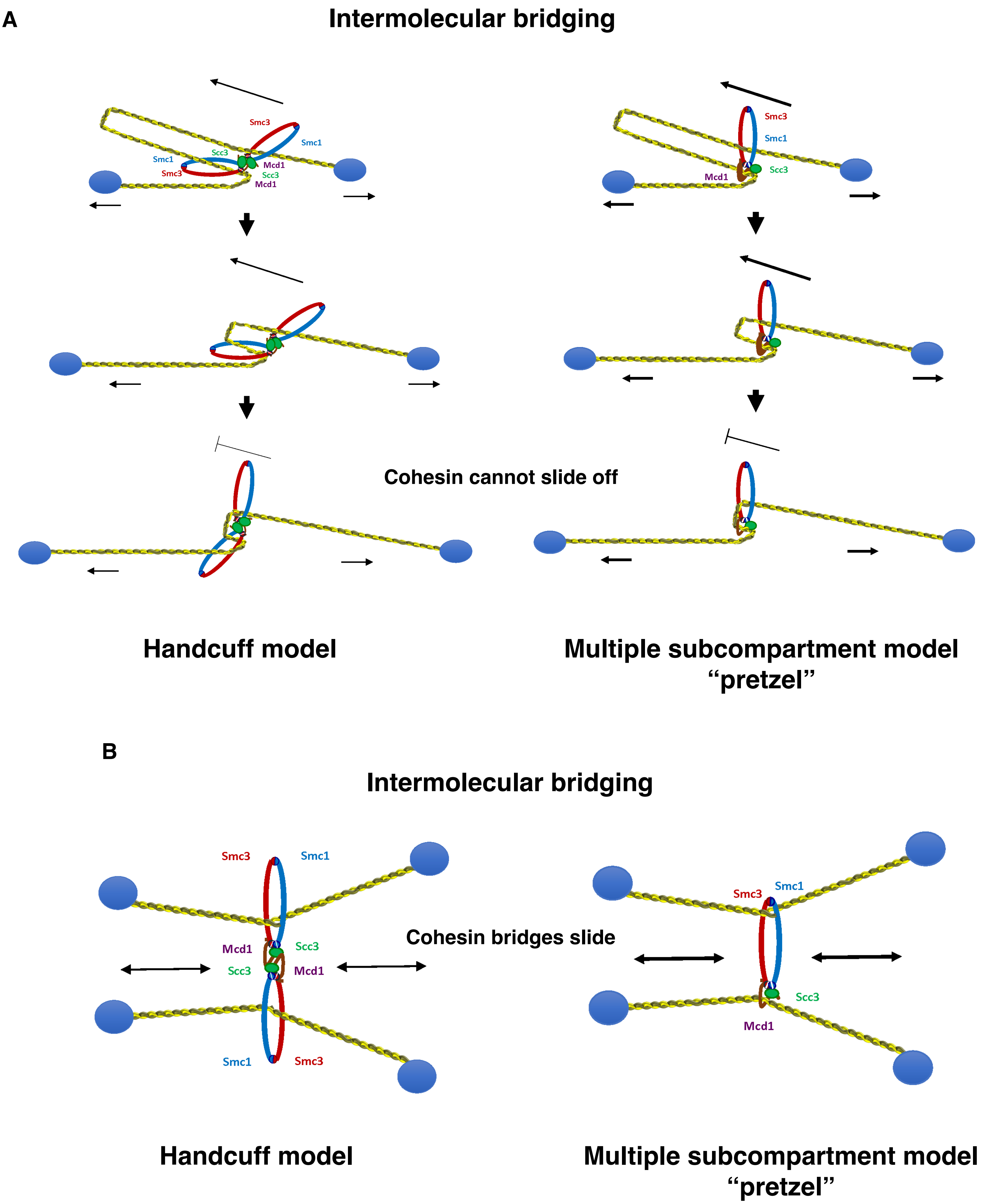
